# Hierarchical chromatin regulation during blood formation uncovered by single-cell sortChIC

**DOI:** 10.1101/2021.04.26.440606

**Authors:** Peter Zeller, Jake Yeung, Buys Anton de Barbanson, Helena Viñas Gaza, Maria Florescu, Alexander van Oudenaarden

**Affiliations:** Oncode Institute, Hubrecht Institute-KNAW (Royal Netherlands Academy of Arts and Sciences) and University Medical Center Utrecht, 3584 CT, Utrecht, The Netherlands

## Abstract

Post-translational histone modifications modulate chromatin packing to regulate gene expression. How chromatin states, at euchromatic and heterochromatic regions, underlie cell fate decisions in single cells is relatively unexplored. We develop sort assisted single-cell chromatin immunocleavage (sortChIC) and map active (H3K4me1 and H3K4me3) and repressive (H3K27me3 and H3K9me3) histone modifications in hematopoietic stem and progenitor cells (HSPCs), and mature blood cells in the mouse bone marrow. During differentiation, HSPCs acquire distinct active chromatin states that depend on the specific cell fate, mediated by cell type-specifying transcription factors. By contrast, most regions that gain or lose repressive marks during differentiation do so independent of cell fate. Joint profiling of H3K4me1 and H3K9me3 demonstrates that cell types within the myeloid lineage have distinct active chromatin but share similar myeloid-specific heterochromatin-repressed states. This suggests hierarchical chromatin regulation during hematopoiesis: heterochromatin dynamics define differentiation trajectories and lineages, while euchromatin dynamics establish cell types within lineages.

## INTRODUCTION

Hematopoietic stem cells (HSCs) reside in the bone marrow (BM) to replenish blood cells while maintaining a balance of diverse blood cell types (Orkin and Zon, 2008; Spangrude et al., 1988). During differentiation, HSCs progressively restrict their potential to fewer lineages to yield mature blood cells (Orkin, 2000). These cell fate decisions are accompanied by gene expression dynamics, which have recently been dissected through single-cell mRNA sequencing technologies (Baccin et al., 2020; Giladi et al., 2018; Paul et al., 2015).

The regulation of gene expression relies, in part, on post-translational modifications of histones that modulate the packing of chromatin (Allfrey et al., 1964; Bell et al., 2011). Chromatin dynamics during hematopoiesis have so far been analyzed in detail for open chromatin regions in single cells (Buenrostro et al., 2018; Ranzoni et al., 2021) and active chromatin marks in sorted blood cell types (Lara-Astiaso et al., 2014). Although the role of repressive chromatin has been characterized in embryonic stem cell cultures (Boyer et al., 2006; Mikkelsen et al., 2007; Peters et al., 2001; Wen et al., 2009) and during early development (Feldman et al., 2006; Fu et al., 2020; Nicetto and Zaret, 2019), the dynamics of repressive chromatin states during hematopoiesis has been relatively unexplored.

Two repressive chromatin states play a major role in gene regulation: a polycomb-repressed state, marked by H3K27me3 at gene-rich, GC-rich regions (Beisel and Paro, 2011; Pauler et al., 2009), and a condensed heterochromatin state mainly found in gene poor AT-rich regions, marked by H3K9me3 (Nicetto and Zaret, 2019). Conventional techniques to detect these histone modifications involve chromatin immunoprecipitation (ChIP), which relies on physical pull-down of histone-DNA complexes. This pull-down can hinder sensitive detection in single cells, although microfluidic extensions of ChIP-seq has improved the assay to single-cell resolution (Grosselin et al., 2019; Rotem et al., 2015). Alternatives to ChIP (Schmid et al., 2004) circumvent this pulldown by using antibody tethering of either protein-A-micrococcal nuclease (pA-MNase) (Ku et al., 2019; Skene and Henikoff, 2017) or protein-A-Tn5 transposase (Harada et al., 2019; Kaya-Okur et al., 2019), improving signal-to-noise by cutting only at specific sites of the genome. Although these tethering-based strategies have enabled large-scale profiling of histone modifications in single cells (Bartosovic et al., 2021; Janssens et al., 2020; Wu et al., 2021), they generally do not enrich for different cell types, making it difficult to dissect chromatin regulation in rare cell types, such as hematopoietic stem and progenitor cells in the bone marrow.

Here we develop sortChIC, which combines single-cell histone modification profiling with cell enrichment, and apply it to map histone modifications in hematopoietic stem cells/early progenitors (HSPCs) and mature blood cell types in the mouse bone marrow. We characterize active (H3K4me1 and H3K4me3) and repressive chromatin (H3K27me3 and H3K9me3). Active chromatin in HSPCs primes for different blood cell fates, while H3K27me3 repressive chromatin in mature cell types silences genes of alternative fates. Although H3K27me3 and H3K9me3 repressive modifications target distinct regions of the genome, most regions that gain or lose repressive modifications during differentiation do so independent of the specific cell fate. By contrast, active chromatin shows divergent changes during hematopoiesis, where gains and losses at genomic regions depend on the specific cell fate. Transcription factor (TF) motif analysis predicts cell type-specifying TFs that drive different chromatin dynamics to acquire distinct chromatin states. Simultaneous targeting of H3K4me1 and H3K9me3 reveals that cell types within the myeloid lineage have distinct active chromatin states, while sharing similar lineage-specific heterochromatin. Our resource reveals a single-cell view of chromatin state dynamics during hematopoiesis in both euchromatic and heterochromatic regions. We propose a hierarchical differentiation program of chromatin regulation in hematopoiesis, by which heterochromatin states define a differentiation trajectory and lineages, while euchromatin states establish cell types.

## RESULTS

### SortChIC maps histone modifications in single cells

To detect histone modifications in single cells, we first fix cells in ethanol to preserve surface antigens for FACS sorting and incubate with an antibody against a particular histone modification. We then add protein A-MNase (pA-MN) which binds to the antibody at specific regions of the genome (Figure 1A). During this incubation, MNase is kept inactive (i.e., no Ca^2+^). After washing away unbound antibody, single cells in the G1 phase of the cell cycle are sorted into 384-well plates (Figure S1A). Next, MNase is activated by adding calcium, allowing MNase to digest internucleosomal regions of the DNA that are in proximity to the antibody. Without the need for purification steps, nucleosomes are stripped off the DNA, and the genomic fragments are ligated to barcoded adapters containing a unique molecular identifier (UMI) and cell-specific barcode. The genomic fragments are amplified by *in vitro* transcription and PCR, and sequenced (Methods).

**Figure 1.**
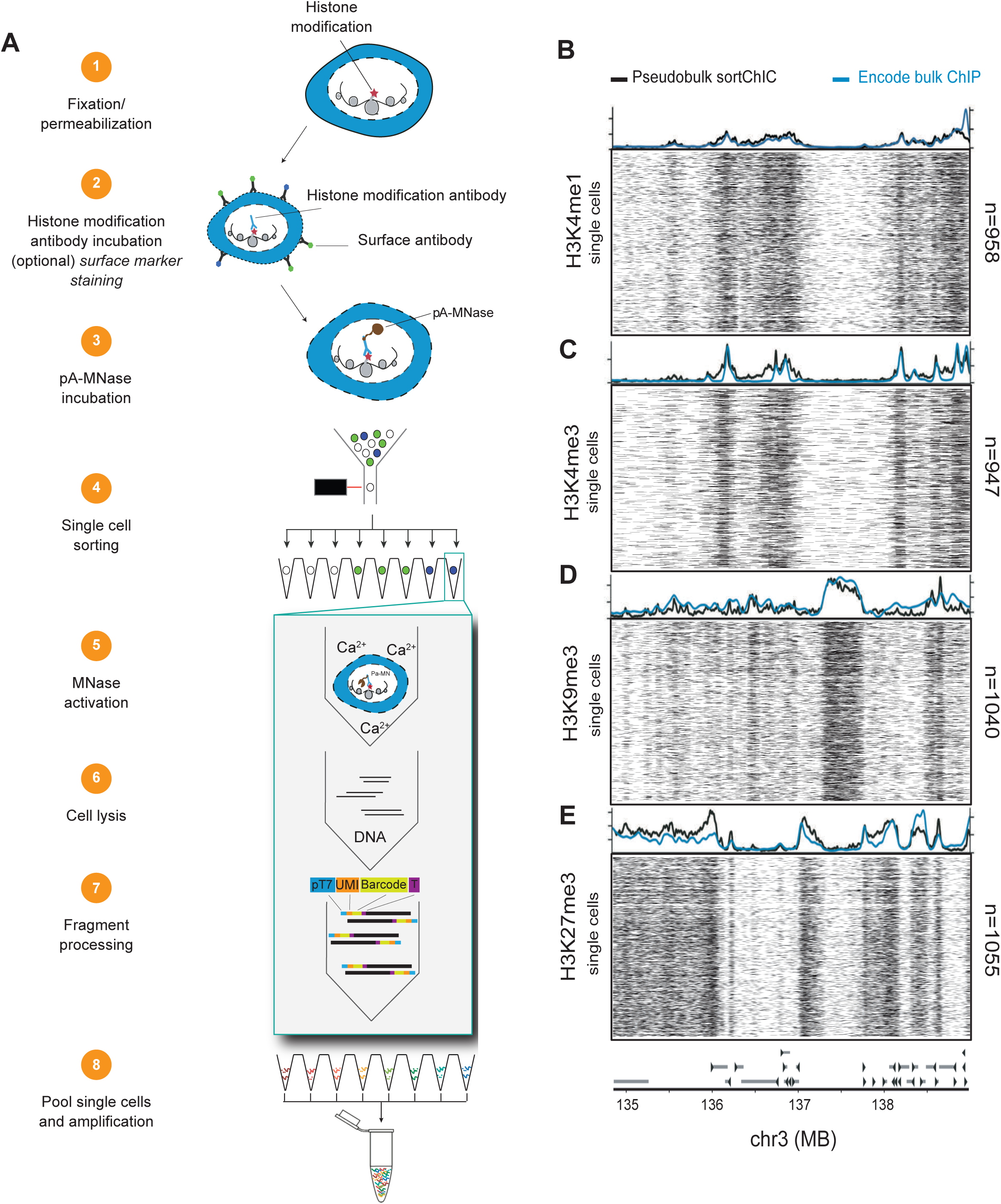
sortChIC maps histone modifications in single cell. (A) Schematic of the sortChIC method. Fixed and permeabilized cells are stained with an antibody targeting a histone modification. Inactive protein A-micrococcal nuclease (pA-MNase) is added, tethering MNase to the histone modification antibody. Single cells are FACS sorted. MNase is activated to induce specific cuts in the genome. Unique molecular identifiers (UMI) and cell-specific barcodes are ligated to the cut fragments. Barcoded fragments are pooled, amplified, and sequenced. (B-E) Location of cuts in H3K4me1 (B), H3K4me3 (C), H3K9me3 (D), and H3K27me3 (E) in individual K562 cells along a 4 MB region of chromosome 3. Black traces represent the sortChIC signal averaged over all individual cells, blue traces represent ENCODE ChIP-seq profiles.

To test if sortChIC is sensitive enough to detect histone modifications in single cells, we apply it on the well-characterized human leukemia cell line K562. In these cells, we map four histone modifications that represent major chromatin states regulating gene expression (Figure 1B-E). For modifications associated with gene activation, we profile H3K4me1 (Figure 1B), found at active enhancers and promoters and H3K4me3 (Figure 1C), found at the promoters of active genes (Heintzman et al., 2007). For modifications associated with repression, we profile H3K9me3 (Figure 1D) and H3K27me3 (Figure 1E), found in gene-poor and gene-rich regions, respectively (Pauler et al., 2009).

For each of the four histone modifications, we process 1128 K562 cells in the G1 phase of the cell cycle to ensure a single genome copy per cell. Using the position of the MNase cut site and unique molecular identifies (UMIs), we map unique MNase cut sites genome-wide (Methods). We use a combination of total unique cuts recovered and fraction of cuts in the MNase-preferred AT context to remove low quality cells (Figure S1B, Methods). Overall, 4176 / 4608 of the cells met our criteria, with a mean of 15000 unique cuts per cell (Figure S1B) with a median of 80% of the sequenced reads falling in peaks identified by adding the sortChIC signal over all cells (Figure S1C, Methods).

We compare pseudobulk sortChIC profiles with publicly available bulk ChIP-seq results (Davis et al., 2018), showing high correlation (Pearson correlation > 0.8) between sortChIC pseudobulk and the ChIP-seq signal for each of their respective marks (Figure S1D-E). Single-cell tracks underneath each average track (Figure 1B) illustrate the high reproducibility of the signal between cells. Of note, the H3K9me3 histone modification profiles obtained from sortChIC represent the heterochromatin state without the need for input normalization (Figure S1F), a procedure that is often needed in classical ChIP experiments (Teytelman et al., 2009). Overall, sortChIC accurately reveals active and repressive chromatin landscapes in single cells.

### Active chromatin in HSPCs primes for different blood cell fates, while H3K27me3 repressive chromatin in differentiated cell types silences genes of alternative fates

We next map active and repressive chromatin changes during blood formation. We combine sortChIC with cell surface marker staining against lineage markers, Sca-1, and c-Kit to sort abundant and rare cell types from the mouse bone marrow in parallel and map histone modifications associated with different chromatin states (Figure S2A). Dimensionality reduction based on a multinomial model (Methods) and then visualizing this latent space with Uniform Manifold Approximation and Projection (UMAP) reveals distinct clusters that contain LSKs (Lin^-^Sca1^+^cKit^+^ sorted cells), unenriched cell types, and mixtures of lineage negative (Lin^-^) and unenriched cell types (Figure 2A, Figure S2B). We use the H3K4me3 signal in cluster-specific promotor regions (transcription start site (TSS) +/− 5 kb) to determine marker genes for eight blood cell types (Figure 2B, Methods). These regions are associated with known cell type-specific genes such as the B cell-specific transcription factor, *Ebf1* (Figure 2C), and the neutrophil-specific gene, *S100a8* (Figure 2D). For H3K4me1 and H3K4me3, these regions are marked in their respective cell types, while for H3K27me3 these regions show specific depletion in their respective cell types (Figure 2E). Using a publicly available dataset (Giladi et al., 2018), we analyze the mRNA abundances associated with our cell type-specific regions across blood cell types and confirm that these sets of genes are cell type-specific (Figure S2C). Our sortChIC data produces high resolution maps of histone modifications in single cells. For example, the TSS of a B cell-specific transcription factor, *Ebf1*, shows B-cell specific signal in H3K4me1 and H3K4me3. For H3K27me3, *Ebf1* is upregulated in non-B cells and depleted in B cells (Figure 2F and Figure S2D-F). Interestingly, we find that hematopoietic stem and early progenitor cells (HSPCs) already have H3K4me3 and H3K4me1 marks at the *Ebf1* promoter and gene body, respectively, suggesting HSPCs may already have active marks at intermediate levels relative to differentiated cell types.

**Figure 2.**
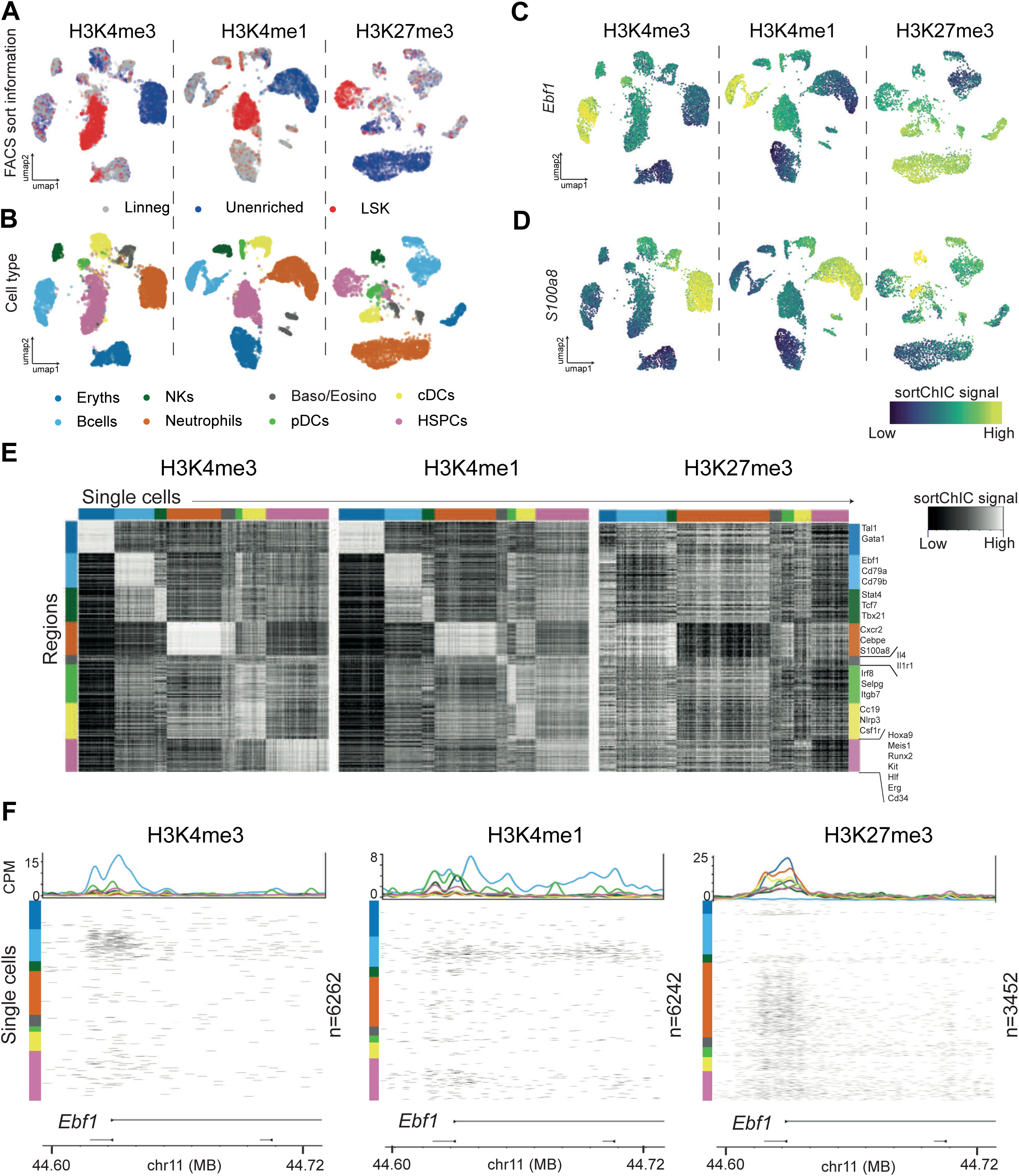
Active and repressive chromatin states in single cells from the mouse bone marrow. (A) UMAPs of H3K4me3, H3K4me1, and H3K27me3 single-cell epigenomes from whole bone marrow (unenriched), lineage negative (Lin^-^), and Lin-Sca1^+^cKit^+^ (LSK) sorted populations. (B) UMAPs colored by cell type. Eryths: erythroblasts, NKs: natural killer cells, Baso/Eosino: basophils/eosinophils, pDCs: plasmacytoid dendritic cells, cDCs: common dendritic cells, HSPCs: hematopoietic stem cells and early progenitor cells, (C) UMAP summary colored by sortChIC signal in a region +/− 5 kb centered at the transcription start site of *Ebf1*, a B cell-specific gene. (D) Same as (C) but for a region around *S100a8*, a neutrophil-specific gene. (E) Heatmap of sortChIC signals for regions around cell type-specific genes showing high levels of active marks (H3K4me1, H3K4me3) in their respective cell type, and correspondingly low levels in the repressive mark (H3K27me3). (F) Example of active and repressive chromatin states near the transcription start site of a B cell specific transcription factor *Ebf1*. H3K4me3 and H3K4me1 show large number of cuts specifically in B cells; H3K27me3 shows B cell-specific depletion of cuts. Colored line plots (same color code as in B) represent the average sortChIC signal for cells of the same cell. Individual cells are ordered by cell type, color coded on the left.

We extend the *Ebf1* observation to all TSSs in our cell type-specific gene sets. To quantify the changes as HSPCs differentiate into different cell types, we compared fold changes between differentiated cell types relative to HSPCs across each of our eight sets of cell type-specific genes derived from H3K4me3 (Figure S2D-F, Methods). When compared to the HSPCs, we find that changes in active chromatin levels are up- or down-regulated depending on the cell fate. For example, at B cell-specific genes, active chromatin levels increase from HSPCs to B cells and to pDCs, but decrease in basophils/eosinophils, neutrophils, and erythroblasts (Figure S2D, E). This divergent pattern occurs in all differentiated cell type-specific gene sets, suggesting that cell type-specific regions in HSPCs already have an intermediate level of active chromatin marks, which are then modulated up or down depending on the cell type into which the HSPCs differentiate.

Repressive H3K27me3 marks at B cell-specific genes, by contrast, are upregulated in non-B cells compared to HSPCs, while only a subset of them loses H3K27me3 when differentiating into B-cells (Figure S2F). Across other cell type-specific genes, we observe a similar trend where HSPCs upregulate H3K27me3 at genes specific for alternative cell fates, likely silencing cell type-inappropriate genes. This upregulation suggests that a main role of H3K27me3 during hematopoiesis is to silence genes of alternative blood cell fates.

In sum, our analysis at blood cell type-specific genes shows that active chromatin primes HSPCs for different blood cell fates, while H3K27me3 repressive chromatin during hematopoiesis silences genes of alternative fates.

### Dynamic H3K9me3 regions reveal clusters enriched for HSPCs, erythroid, myeloid, and lymphoid lineages

To understand chromatin regulation in gene-poor heterochromatic regions, we map H3K9me3 modifications from the same technical batch of HSPCs, lineage negative, and unenriched bone marrow cells as was used for the H3K4me1, H3K4me3, and H3K27me3 sortChIC experiments. In contrast to the other three marks, where we find eight cell types, H3K9me3 analysis reveals four clusters: one cluster containing mostly LSKs, one cluster containing mostly unenriched cells, and two clusters containing a mixture of unenriched and lineage negative cells (Figure 3A, B). We find large megabase-scale domains marked by H3K9me3 in gene-poor regions that are constant across cell types, but also smaller sub-megabase regions with cluster-specific signal, suggesting that there may be differences reflecting different cell types or lineages, despite having large regions of heterochromatin that are conserved across cell types (Figure 3C). Differential analysis on 50 kb regions across the genome identified 6085 cluster-specific regions (q-value < 10^−9^, deviance goodness-of-fit test from Poisson regression, Methods) for H3K9me3. These cluster-specific H3K9me3 regions have a median distance of 62.8 kb to the nearest TSS of a gene, and are closer to TSSs than H3K9me3-marked regions in general, which have a median distance of 138 kb to a TSS (Figure S3A). This suggests that some cluster-specific H3K9me3 regions may be associated with gene regulation.

**Figure 3.**
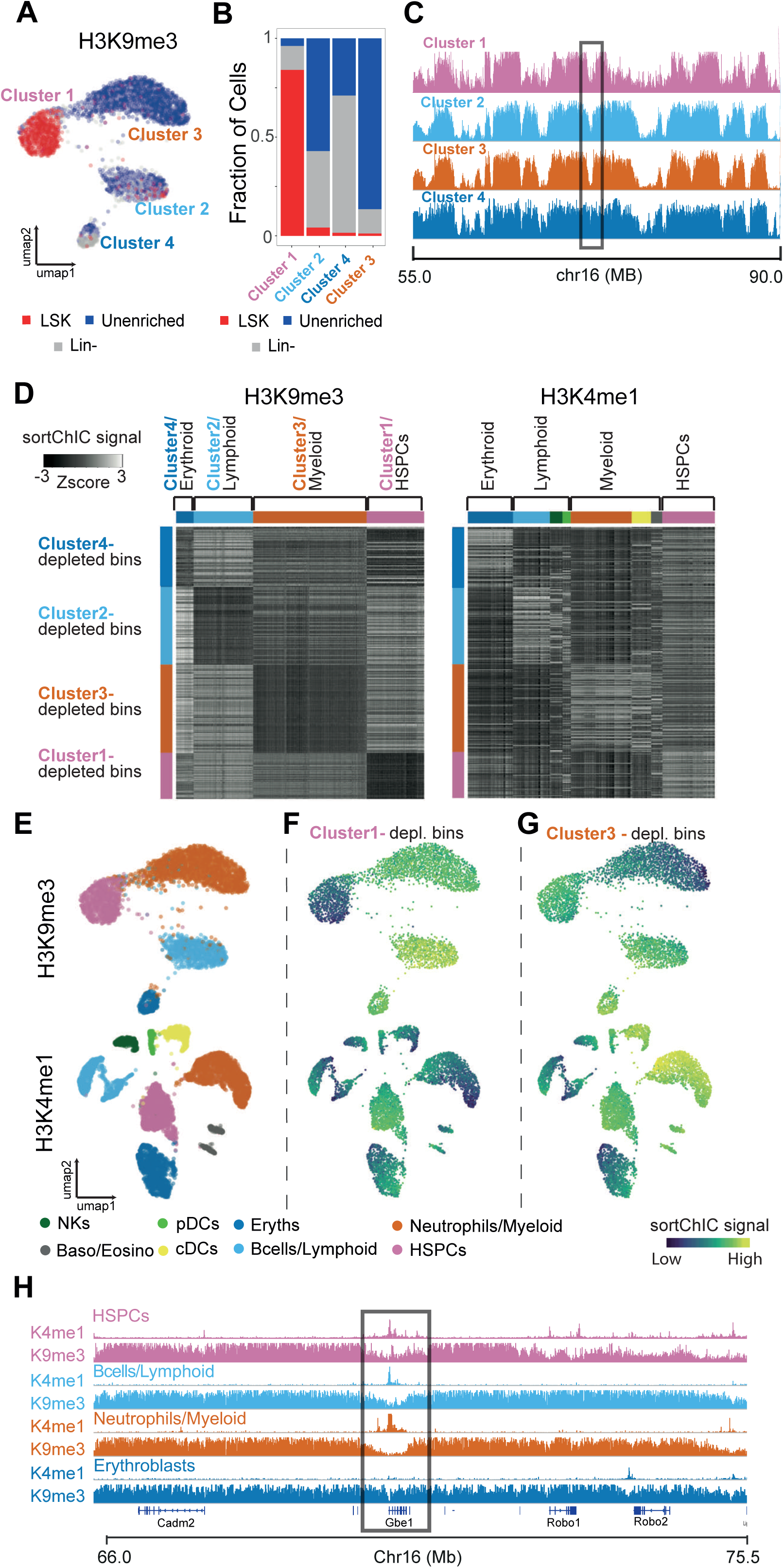
Heterochromatin state dynamics during hematopoiesis. (A) UMAP of H3K9me3 representing single cells from whole bone marrow (unenriched), lineage negative (Lin-), and Lin-Sca1+cKit+ (LSK) sorted cells. (B) Fraction of unenriched, Lin^-^, and Lin^-^Sca1^+^cKit^+^ cells in each of the four H3K9me3 clusters. (C) Region showing the H3K9me3 pseudobulk sortChIC signal of the four clusters. (D) Heatmap of 50 kb bins displaying the relative H3K9me3 (left) and H3K27me3 (right) sortChiC signal in erythroblasts, lymphoid, myeloid, and HSPCs. (E) UMAP of H3K4me1 and H3K9me3 sortChIC data, colored by cell type. (F) Single-cell signal of cluster1-depleted bins (averaged across the 150 bins) showing low H3K9me3 and high H3K4me1 signal in lymphoid cells. Both H3K9me3 (above) and H3K4me1 (below) are quantified using the same set of bins. (G) Single-cell signal of cluster3-specific bins showing low H3K9me3 and high H3K4me1 signal in myeloid cells. (H) Zoom-in of the same genomic region in (B) for H3K9me3 and H3K4me1 pseudobulk sortChIC signal.

Since cluster-specific H3K9me3 regions were closer to TSSs than general H3K9me3-marked regions, we hypothesize that the H3K4me1 mark in these same regions may also show cluster-specific signal. Out of the 6085 cluster-specific H3K9me3 regions, we select 150 regions with the largest depletion of the H3K9me3 sortChIC signal relative to HSPCs for each of the four clusters, resulting in four sets of cluster-specific regions (Figure S3B). Comparing the H3K4me1 signal in each of these four sets of regions shows cell type-specific retention in these regions (Figure S3C), consistent with an anticorrelated relationship with H3K9me3. Heatmaps of the H3K9me3 and H3K4me1 signal at the four sets of regions reveal an anticorrelated structure between H3K9me3 versus H3K4me1 (Figure 3D), allowing the prediction of cell types in the H3K9me3 data (Figure S3B, C). Erythroid regions are upregulated in erythroblasts in H3K4me1; lymphoid regions show strongest upregulation in B cells; myeloid regions show strongest upregulation in neutrophils. We use this anticorrelation at cluster-specific H3K9me3 regions to identify cells related to erythroid, lymphoid, and myeloid lineages in H3K9me3 (Figure 3E). We find that regions depleted of H3K9me3 in HSPCs show upregulation of H3K4me1 in HSPCs (Figure 3F). For H3K9me3-depleted regions in myeloid cells, we find that H3K4me1 is upregulated not only in neutrophils, but also in other cell types that share the myeloid lineage, such as common dendritic cells (Figure 3G). This anticorrelation is exemplified in a genomic region surrounding the *Gbe1* gene, showing repression of H3K4me1 signal specifically in erythroblasts accompanied by high levels of H3K9me3. In this region, HSPCs, lymphoid, and myeloid cell types show enrichment of H3K4me1 accompanied by a marked depletion in H3K9me3 (Figure 3H). At these lineage-specific H3K9me3 regions, we also see cell type-specific signal in H3K4me3 and in H3K27me3, although the pattern is weaker than in H3K4me1 (Figure S3D). Overall, we find the H3K9me3 clusters are related to HSPCs, erythroid, lymphoid, and myeloid lineages.

### Repressive chromatin dynamics are largely cell fate-independent

We ask whether global patterns in chromatin dynamics during hematopoiesis differed between repressive and active marks. We apply differential analysis on 50 kb regions for all four marks, resulting in 10518 dynamic bins for H3K4me1, 2225 for H3K4me3, 5494 for H3K27me3, and 6085 for H3K9me3 (q-value < 10^−50^ for H3K27me3, H3K4me1, and H3K4me3; q-value < 10^−9^ for H3K9me3, Table S1, Methods). For each histone modification, we cluster the pseudobulk signal of each cell type across the bins. Hierarchical clustering reveals global relationships between cell types. In active marks, we find that the largest differences come from erythroblast versus non-erythroblasts (Figure 4A, left two panels). This erythroblast distinction corroborates with our analysis of fold changes at TSSs of cell type-specific genes, where the erythroblasts shows the largest changes in active chromatin during differentiation (Figure S2D, E). Furthermore, we find that HSPCs often have intermediate levels of H3K4me1 and H3K4me3 (Figure 4A, left two panels), suggesting a generally more accessible chromatin state HSPCs.

**Figure 4.**
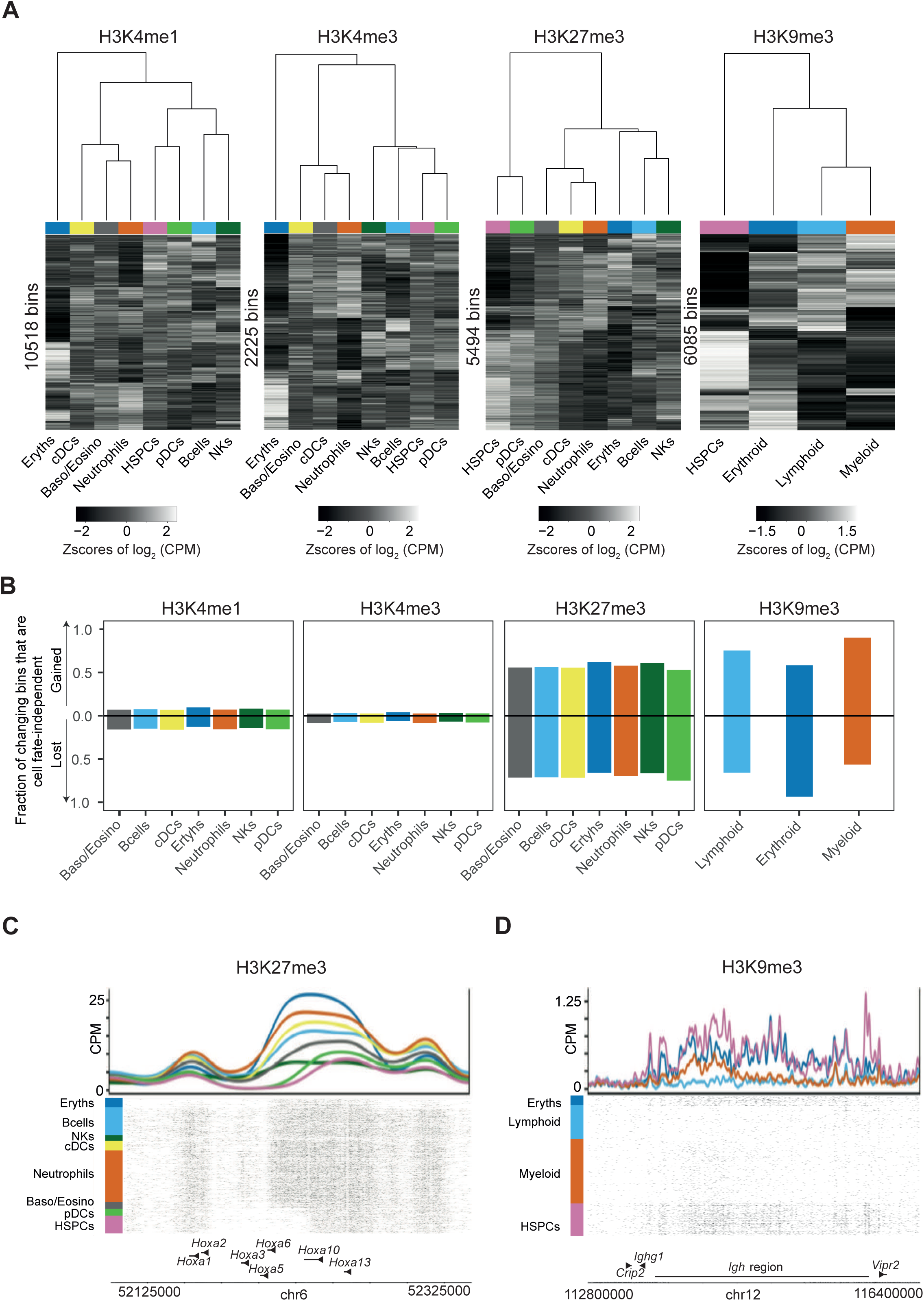
Repressive chromatin dynamics are largely cell fate-independent. (A) Heatmap of log_2_ counts per million (CPM) of 50 kilobase bins across pseudobulks. Changing bins that are statistically significant are shown (deviance goodness-of-fit test from Poisson regression, Methods). The rows and columns are ordered by complete-linkage clustering. Above each heatmap is a dendrogram from clustering the columns, showing the relationship between cell types. (B) Barplot of the fraction of changing bins (Methods) that are gained or lost in all non-HSPCs relative to HSPCs. Each cell type shows two bars, one for each direction (either gained or lost). Fraction is calculated by dividing the number of bins that change cell fate-independently by the number of bins that change in that cell type for that direction. (C) Genome browser view of the *Hoxa* region showing a H3K27me3 domain that is gained during hematopoiesis. (D) Genome view of the immunoglobulin heavy chain (*IgH*) region displaying the loss of a H3K9me3 domain in lymphoid and myeloid cells.

Projecting the active mark data onto the two most significant axes of chromatin variation (Townes et al., 2019), shows that the HSPCs take a central position relative to other cell types, suggesting that changes in active chromatin during hematopoiesis can diverge depending on the specific cell fate (Figure S4A, left two panels).

By contrast, repressive chromatin dynamics, marked by H3K27me3 and H3K9me3, mainly distinguish between HSPCs and differentiated cell types, thereby marking the progress along the differentiation trajectory (Figure 4A, right two panels). Projecting the repressive mark data reveals an axis connecting HSPCs and other cell types (Figure S4A, right two panels). To ask whether regions gain or lose chromatin marks depending on the specific cell fate, we calculate the fraction of changing bins that gain or lose chromatin marks in all non-HSPCs relative to HSPCs (Methods). We find more than half of bins that gain or lose repressive marks between one cell type versus HSPCs are also gaining or losing marks across all other cell fates (Figure 4B), suggesting that many changes in repressive chromatin during hematopoiesis occur independent of the specific cell fate. By contrast, only 8 percent of bins in active chromatin, on average, show cell type-independent changes. Fold changes between HSPCs and non-HSPCs at changing bins show distinct separation between HSPCs and non-HSPCs in repressive marks, but not in active marks (Figure S4B), corroborating that many changes in repressive chromatin are independent of cell fate. These cell fate-independent changes are exemplified for H3K27me3 at the *Hoxa* region, which shows low levels of H3K27me3, and its levels are upregulated in differentiated cell types (Figure 4C). HSPCs at the *Igh* region show high levels of H3K9me3, and its levels are downregulated in myeloid and lymphoid cells, suggesting that this *Igh* region, which encodes the heavy chains of immunoglobulins, are de-repressed during differentiation (Figure 4D).

We ask whether H3K27me3 and H3K9me3 may regulate distinct processes. We confirm that H3K27me3 dynamics occur at GC-rich regions close to TSSs while H3K9me3 dynamics at AT-rich regions occur further from TSSs (Figure S4C, D), consistent with known sequence-specific contexts of the two repressive marks (Pauler et al., 2009). GO term analysis of H3K9me3 regions unique to HSPCs shows enrichment for immune-related processes such as phagocytosis, complement activation, and B cell receptor signaling (Figure S4E), suggesting that HSPCs use H3K9me3 to repress genes that may later need to be used in differentiated blood cells. By contrast, GO term analysis of H3K27me3 regions unique to HSPCs does not show consistent enrichment for biological processes related to blood development.

Taken together, we find that HSPCs have active chromatin marks at intermediate levels relative to other blood cell types. During differentiation, active marks at these regions can then be up- or down-regulated depending on the specific cell fate. By contrast, most dynamic repressive chromatin regions are gained or lost independent of the specific cell fate.

### Transcription factor motifs underlie active and repressive chromatin dynamics in hematopoiesis

Next, we ask whether regulatory information in the DNA sequences underlying the sortChIC data can explain the cell type-specific distributions in active and repressive chromatin landscapes. We hypothesize that regions with correlated sortChIC signal across cells can be explained in part by transcription factor binding motifs shared across these regions (Arnold et al., 2012, 2013) (Figure S5A, Methods). Overlaying the predicted single-cell TF motif activities onto the UMAP representation shows the expected blood cell type-specific activity for known regulators. We find the ERG motif active specifically in HSPCs (Figure 5A, left), consistent with the role of ERG in maintenance of hematopoietic stem cells (Knudsen et al., 2015). CEBP family motif is active in neutrophils (Figure 5A, mid left), consistent with its proposed function (Cloutier et al., 2009; Scott et al., 1992). EBF motif activity is specific to B cells (Figure 5A, mid right), consistent with its role in specifying B cell differentiation (Vilagos et al., 2012). We find TAL1 to have erythroblast-specific activity (Figure 5A, right), in agreement with its role in erythropoiesis (Hall et al., 2005).

**Figure 5.**
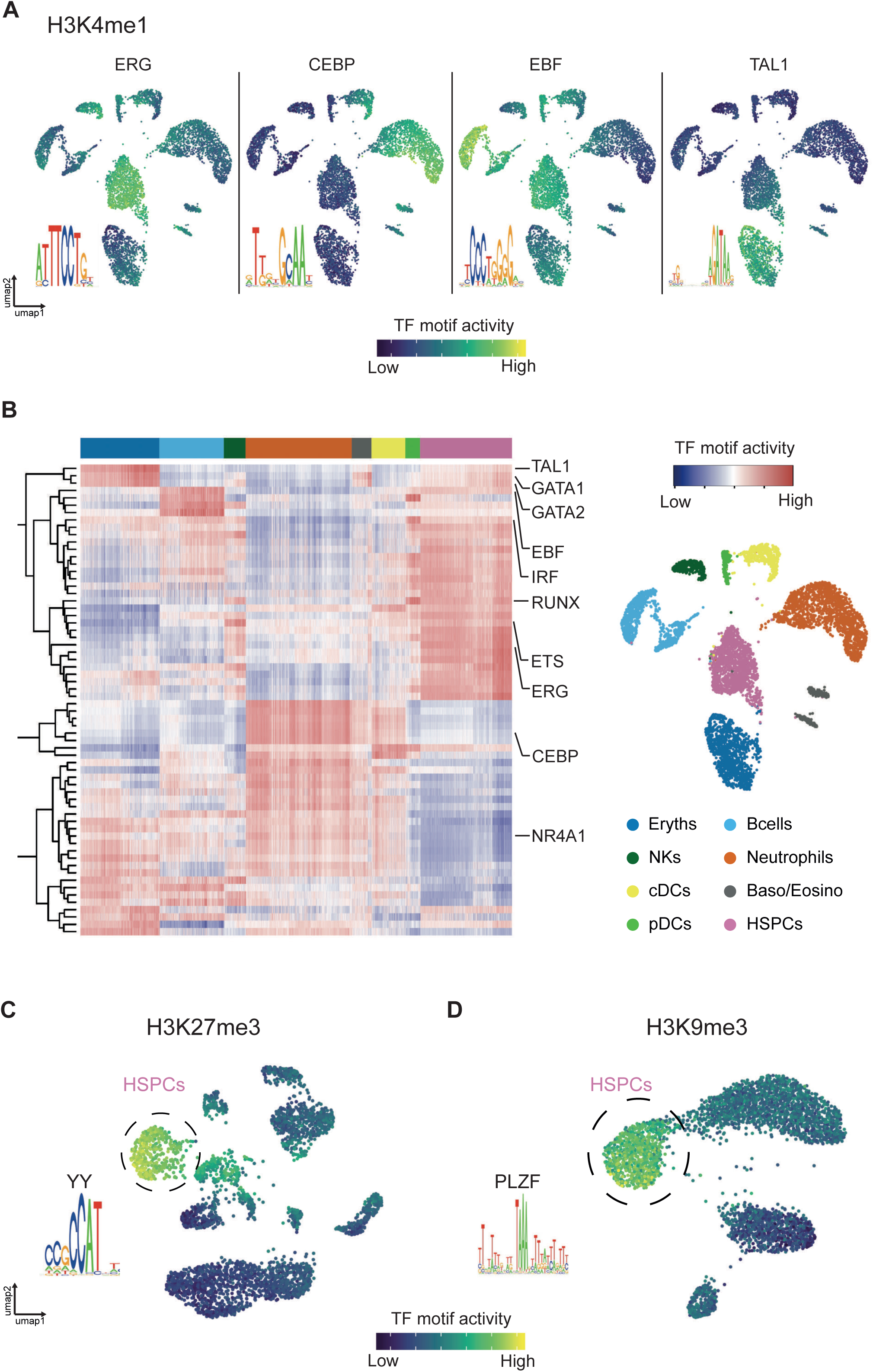
Transcription factor motifs underlie active and repressive chromatin dynamics in hematopoiesis. (A) Examples of four transcription factor (TF) motifs whose activities are predicted to drive cell type-specific H3K4me1 distributions. The ERG motif is predicted to be active in HSPCs, the CEBP motif in neutrophils, the EBF motif in B cells, and the TAL1 motif in erythroblasts. Cell type for each cell cluster is labeled in (B). (B) Heatmap of H3K4me1 TF motif activities in single cells. Rows represent motifs. Columns are individual cells whose cell types are annotated by the top color bar. The right panel shows a H3K4me1 UMAP colored by cell types, with cell type-to-color legend below. (C) Predicted H3K27me3 activity of a motif belonging to the Yin Yang (YY) protein family in single cells. Circled cluster is enriched for HSPCs. (D) Predicted H3K9me3 activity of PLZF motif in single cells. Circled cluster is enriched for HSPCs.

We summarize the inferred single-cell TF activities in a heatmap to comprehensively predict transcriptional regulators underlying the cell type-specific distribution of active chromatin (Figure 5A, B). We predict motifs active in pDCs belonging to the IRF and RUNX family (Figure 5B, Figure S5B-D). High activity of IRF motifs corroborates with pDC function to secrete type 1 interferon (Honda et al., 2005; Wang et al., 2016). Runx proteins have been shown to regulate development of dendritic cell progenitors (Satpathy et al., 2014) and migration of pDCs from the bone marrow (Sawai et al., 2013). We find NK cells to have high ETS family motif activity (Figure 5B, Figure S5B, E), consistent with the role *of Ets1* in development of natural killer and innate lymphocyte cells (Barton et al., 1998; Zook et al., 2016). Finally, we predict transcription factors that have low activity in HSPCs and pDCs but high activity in other cell types, such as the NR4A family (Figure 5B, Figure S5, B and F*). Nr4a1* has been shown to repress gene expression (Nowyhed et al., 2015) and control hematopoietic stem cell quiescence by suppressing inflammatory signaling (Freire and Conneely, 2018). The low activity of several TFs specific in HSPCs and pDCs suggests that the pDCs we identify could be in a more progenitor-like state, consistent with the pseudobulk clustering results in H3K4me1, H3K4me3 and H3K27me3 (Figure 4A).

We apply our TF motif analysis to the two repressive chromatin landscapes to predict motifs that explain HSPC-specific distributions of repressive chromatin. In H3K27me3, we predict a CCAT motif belonging to the Yin Yang family (Kim et al., 2007) specifically active in HSPCs (Figure 5C). Of note, *Yy1* is gene encoding the polycomb group protein and has been shown to regulate hematopoietic stem cell self-renewal (Lu et al., 2018). In H3K9me3, we predict an AT-rich motif belonging to the transcriptional repressor PLZF specifically active in HSPCs (Figure 5D). PLZF has been implicated in regulating the cell cycle of hematopoietic stem cells (Vincent-Fabert et al., 2016).

In sum, our motif analysis explains differences in chromatin levels across cells in terms of TF activities. Our predictions suggest that differentiating blood cells decide which active regions to up- or down-regulate depending on the cell type-specific TFs that associate with different regions. Although repressive chromatin dynamics in both H3K27me3 and H3K9me3 are mainly cell fate-independent, our analysis suggests that distinct TFs regulate the two separate pathways.

### Distinct cell types can share similar heterochromatin landscapes

To understand in more detail the relationship between the eight cell types identified by histone marks of gene-rich regions (H3K4me1, H3K4me3, and H3K27me3) to the four clusters identified by H3K9me3, we stain cells with both H3K4me1 and H3K9me3 antibody concurrently (Yeung et al., 2021). This double-incubation strategy generates cuts that come from both H3K4me1 and H3K9me3 from the same cell, and uses our sortChIC resource to infer the relationships between the two marks in single cells (Figure 6A). We sort Lin^-^ and unenriched cells to profile both abundant and rare cell types. UMAP of the joint landscape reveals clusters that are depleted of mature lineage markers as well as enriched for mature cell types (Figure 6B). We use clusters from H3K4me1 and H3K9me3 single-incubated data to develop a model of how the double-incubated data could be generated (Figure 6C, Methods).

**Figure 6.**
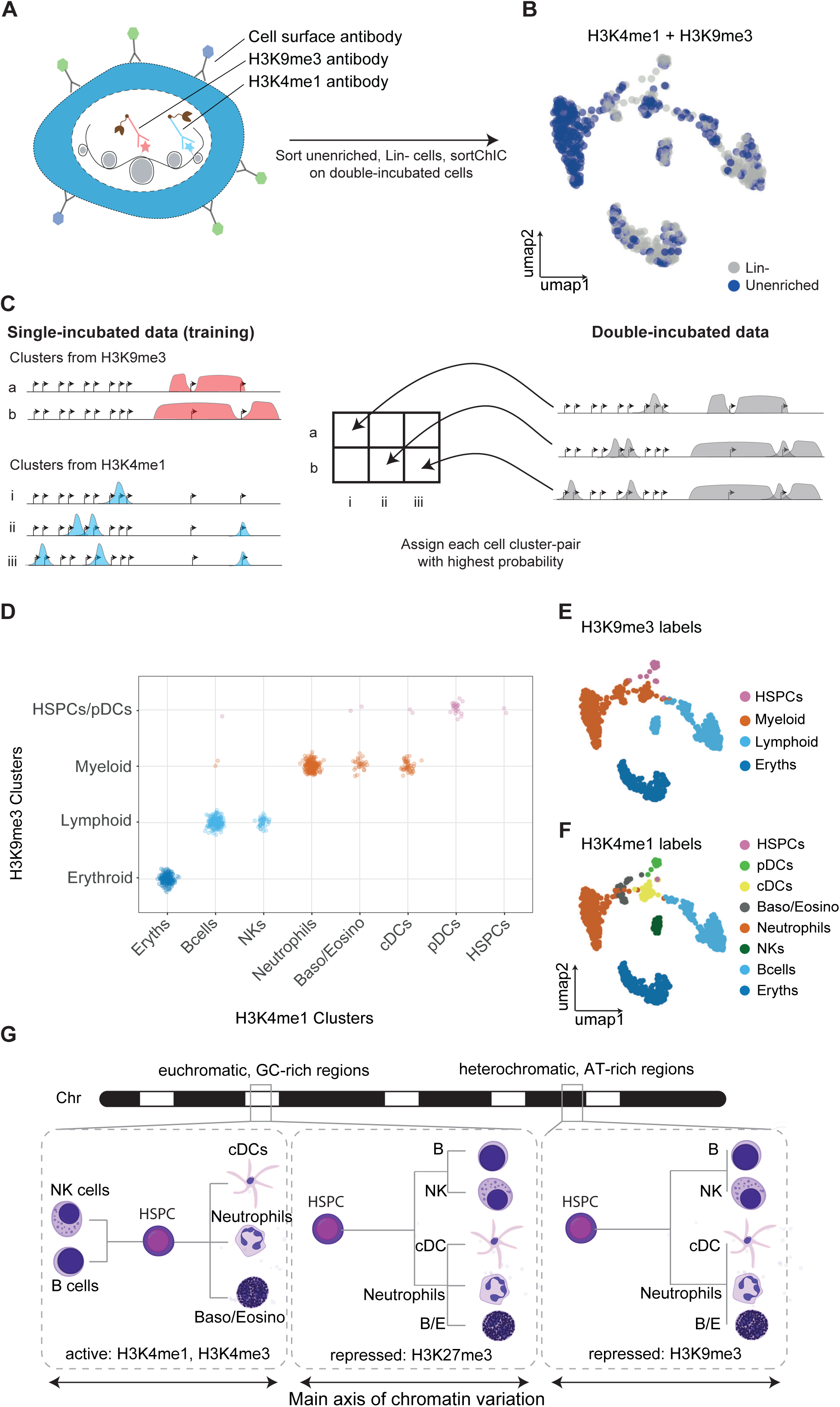
Distinct cell types can share similar heterochromatin landscapes. (A) Double incubation experiment produces cuts associated with either H3K4me1 or H3K9me3. (H3K4me1+H3K9me3) (B) UMAP representation of the H3K4me1+H3K9me3 landscape in unenriched and lineage-negative cells in the bone marrow. (C) Schematic of how the standard single-incubated data can produce a model of which cluster-pair (one from H3K9me3, the other from H3K4me1) generates the observed double-incubated data. (D) Output of cluster-pair predictions from H3K4me1+H3K9me3 double-incubated cells. Cells are colored by their predicted H3K9me3 clusters. (E and F) UMAP representation of the H3K4me1+H3K9me3 landscape, colored by their predicted H3K4me1 cluster (E) or H3K9me3 cluster (F). (G) Graphical summary of chromatin dynamics as dendrograms showing relationships between HSPCs and differentiated cells. During hematopoiesis, the direction of change in active chromatin depends on the specific cell fate, resulting in global differences that are largest between differentiated cell types from different lineages. By contrast, many regions gain or lose repressive marks during hematopoiesis independent of the specific cell fate, resulting in global differences that are largest between HSPCs and differentiated cell types. Dynamics in active marks and H3K27me3-marked repressive chromatin reveal cell type information, while dynamics in heterochromatin regions marked by H3K9me3 reveal lineage information.

For our model, we focus on regions that show large differences between cell types. We select 811 regions associated with cell type-specific genes (Methods) found in our H3K4me1 analysis (Figure 2E) and 6085 cluster-specific regions (50 kb bins) found in our H3K9me3 analysis (Figure 4A, right panel) as features in our model, making a total of 6896 regions. To verify that our features show cluster-specific differences, we cluster the single-incubated H3K4me1 signal across cell types (Figure S6A). We find that neutrophils, basophils/eosinophils, and cDCs cluster together, consistent with their common myeloid lineage (Akashi et al., 2000). B cells and NK cells also cluster together, consistent with their common lymphoid lineage (Kondo et al., 1997). Erythroblasts form a distinct branch from other cell types. Finally, pDCs cluster most closely with HSPCs.

Since *a priori* we do not know which cluster from H3K4me1 pairs with which cluster from H3K9me3, we generate an *in silico* model of all possible pairings (Figure 6C, left). For each double-incubated cell, we then perform model selection to choose the pair with the highest probability (Figure 6C, right, and Figure S6C, D; Methods). This selection reveals that neutrophils, basophils/eosinophils, and cDCs share a common heterochromatin landscape, reflecting their myeloid lineage (Figure 6D). We find B-cells and NK cell share a lymphoid-specific heterochromatin. Erythroblasts do not share a heterochromatin landscape with any other cell type. Although we did not explicitly sort for HSPCs, a small fraction of cells was assigned to both an HSPC-specific active chromatin and HSPC-specific heterochromatin state, reflecting the rarity of HSPCs in the bone marrow. Surprisingly, we find pDCs associated with the HSPC-enriched H3K9me3 landscape, suggesting that these cells that we sorted may have already committed towards a pDC fate through its active chromatin, while its heterochromatin remains undifferentiated.

This joint analysis confirms that distinct cell types in related lineages can share their heterochromatin state (Figure 6E, F), suggesting a hierarchical model where changes in heterochromatin establish lineages and changes in active chromatin define cell types within lineages.

## DISCUSSION

We profile and analyze active and repressive chromatin states in single cells during blood formation, providing a comprehensive map of chromatin regulation at both euchromatic and heterochromatic regions. We find that repressive chromatin shows distinct dynamics compared with active chromatin, demonstrating that profiling repressive chromatin regulation in single cells reveals novel dynamics not captured by profiling active chromatin. Active chromatin primes HSPCs, and is up- or down-regulated depending on the specific cell fate, mediated by cell type-specific transcription factors. Consequently, active chromatin shows divergent changes for different blood cell fates, resulting in global differences in active chromatin that are larger between mature cell types than between HSPCs and mature cell types (Figure 6G, left panel). These active chromatin dynamics likely reflect the dynamics in mRNA abundances (Lara-Astiaso et al., 2014). By contrast, changes in repressive chromatin during hematopoiesis often occur in the same direction (either gained or lost) regardless of the specific cell fate, resulting in global differences in repressive chromatin that are larger between HSPCs and mature cell types than between mature cell types (Figure 6G, middle and right panel). Overall, our results show that single-cell repressive chromatin dynamics provide an orthogonal viewpoint to active chromatin and mRNA dynamics during hematopoiesis.

Technologies to profile histone modifications in single cells by sequencing is still in its infancy, but has the potential to unlock the spectrum of chromatin states in the genome of individual cells. The ideal assay strives to have high sensitivity, high throughput, and robustness in both active and repressive chromatin states. Current techniques to map histone modifications in single cells use one of three approaches: ChIP-based, pA-Tn5-based, and pA-MNase-based. ChIP-based strategies utilize microfluidics systems to reduce reaction volumes and overcome the low sensitivity of ChIP (Grosselin et al., 2019; Rotem et al., 2015). pA-Tn5-based strategies profile histone modifications with high sensitivity and throughput, but requires optimizing experimental conditions to reduce the intrinsic affinity of Tn5 to open chromatin regions, especially when studying repressive chromatin states (Bartosovic et al., 2021; Harada et al., 2019; Janssens et al., 2020; Kaya-Okur et al., 2019; Wang et al., 2019; Wu et al., 2021). pA-MNase-based methods profile histone modifications with high sensitivity, and have robust detection of modifications associated with euchromatic regions as well as heterochromatic regions, but has generally less throughput compared with Tn5-based methods (Hainer et al., 2019; Ku et al., 2019). SortChIC is a unique single-cell method that combines cell enrichment to greatly enhance throughput of rare cells, while achieving high sensitivity and robustness to profile active and repressive (both at GC-rich and AT-rich) chromatin states (Table S2).

This comprehensive profiling of rare progenitors and their multiple cell fates enables new systematic analyses, such as quantifying chromatin dynamics that are cell fate-independent during differentiation. This analysis reveals that cell fate-independent changes during differentiation occur frequently for repressive chromatin, while such changes for active chromatin are rare. Our strategy combines rare progenitor cell enrichment with comprehensive differentiated cell type profiling to allow systematic analysis of chromatin dynamics during differentiation into multiple cell fates.

Our single-cell analysis further expands the role of H3K9me3, which has been classically associated with constitutive types of chromatin (Nicetto and Zaret, 2019). We find that H3K9me3 is a dynamic chromatin modification that regulates different lineages in blood and is rewired as HSPCs differentiate into different blood lineages. Although *in vivo* dynamics in H3K9me3 have been recently reported during early development (Feldman et al., 2006; Fu et al., 2020; Nicetto and Zaret, 2019), our results extend the role of H3K9me3 dynamics to also regulate homeostatic renewal in adult physiology.

Joint profiling analysis demonstrates that cell types from a common lineage can share a similar heterochromatin landscape. Our results suggest that the distinct chromatin dynamics in active chromatin and heterochromatin can reveal the hierarchical relationships between cell types. We find cDCs, neutrophils, and basophils/eosinophils to share a similar myeloid-specific heterochromatin landscape, suggesting that the H3K9me3 mediated heterochromatin can be relatively stable while other chromatin changes further distinguish between distinct cell types. pDCs have been reported to come from lymphoid precursors (Dress et al., 2019), although the exact origin has been debated (Rodrigues et al., 2018). We find pDCs to be distinct from cDCs at both the active and heterochromatin level, although the heterochromatin of pDCs is also distinct from other lymphoid cell types such as B cells or NK cells. One explanation could be that pDCs diverge early from other lymphoid cell types and do not participate in the heterochromatin rewiring that occurs during lymphopoiesis. Overall, we propose a hierarchical chromatin regulation program during hematopoiesis, in which heterochromatin states define a differentiation trajectory and lineages, while euchromatin states establish cell types.

## Acknowledgements

We thank R. van der Linden for assistance during experiments. We thank Amir Giladi for sharing mRNA abundance tables of cell types and critical reading of the manuscript. This work was supported by a European Research Council Advanced grant (ERC-AdG 742225-IntScOmics), Nederlandse Organisatie voor Wetenschappelijk Onderzoek (NWO) TOP award (NWO-CW 714.016.001), the Swiss National Science Foundation, and the Human Frontiers for Science Program. This work is part of the Oncode Institute which is partly financed by the Dutch Cancer Society.

## Contributions

P.Z., J.Y., and A.v.O. designed the project; P.Z. and M.F. developed technique; P.Z. and H.V.G. performed experiments; J.Y. developed and applied the statistical methods. B.A.d.B., M.F., and J.Y. wrote the sortChIC demultiplexing and preprocessing pipeline. J.Y., A.v.O, and P.Z. analyzed the data. P.Z., J.Y. and A.v.O, wrote the manuscript.

## METHODS

### Cell culture

K562 cells (ATCC® CCL-243™) were grown in RPMI 1640 Medium GlutaMAX™, supplemented with 5% FCS, Pen-Strep and non-essential amino acids. After harvesting cells were washed 3 times with room temperature PBS before continuing with the sortChIC protocol.

### Animal experiments

Experimental procedures were approved by the Dier Experimenten Commissie of the Royal Netherlands Academy of Arts and Sciences and performed according to the guidelines. Primary bone marrow cells were harvested from 3-months-old C57BL/6 mice. Femur and Tibia were extracted, the bones ends were cut away to access the bone marrow which was flushed out using a 22G syringe with HBSS (-Ca, -Mg, -phenol red, Gibco 14175053) supplemented with Pen-Strep and 1% FCS. The bone marrow was dissociated and debris was removed by passing it through a 70 μm cell strainer (Corning, 431751). Cells were washed with 25 ml supplemented HBSS before linage marker staining was performed following the instructions of the EasySep™ Mouse Hematopoietic Progenitor Cell Isolation Kit (Stemcell) at half of the recommended concentration of the biotinylated antibodies. This was followed by 30min incubation at 4 °C with a Streptavidin-PE (Biolegend, 1:5000), anti c-kit-APC (Biolegend, 1:800) and anti sca1-PeCy7 (Biolegend, 1:400). After 2 additional washes with HBBS (+PS, +FCS) cells were prepared following the sortChIC protocol for the 4 different histone modifications.

### Pa-MN production

The Pa-MN fusion protein was produced following the methods section in (Schmid et al., 2004). pK19pA-MN was a gift from Ulrich Laemmli (Addgene plasmid # 86973; http://n2t.net/addgene:86973; RRID: Addgene_86973)

### sortChIC-seq experiments

#### Cell preparation: fixation

All steps were performed on ice. Cells were resuspended in 300 μl PBS per 1 million cells in a 15 ml protein low binding falcon tube and 700 μl ethanol (−20 °C precooled) per 1 million cells are added while vertexing cells at middle speed. Cells were fixed for 1 h at −20 °C. After fixation cells were washed twice in 1 ml wash buffer (47.5 ml H_2_O RNAse free, 1 ml 1M HEPES pH 7.5 (Invitrogen), 1.5 ml 5M NaCl, 3.6 μl pure spermidine solution (Sigma Aldrich), 0.05% Tween20, protease inhibitor cocktail (Sigma Aldrich) with 4 μl/ml 0.5 M EDTA). For K562 cells 3 plates were sorted for each modification. For BM we sorted 19, 17, 18, and 17 plates for H3K4me1, H3K4me3, H3K27me3, and H3K9me3, respectively.

#### Cell preparation: nuclei

Cells were washed once in 1 ml wash buffer (47.5 ml H2O RNAse free, 1 ml 1M HEPES pH 7.5 (Invitrogen), 1.5 ml 5M NaCl, 3.6ul pure spermidine solution (Sigma Aldrich), 0.05% Saponin, protease inhibitor cocktail (Sigma Aldrich) with 4 μl/ml 0.5 M EDTA). Nuclei were isolated by further Saponin incubation overnight in parallel to the antibody staining. For BM we sorted 9 plates each for H3K4me1, H3K4me3, and H3K9me3.

#### Antibody staining

Cells were pelleted at 500g for 4 min and resuspended in 200 μl Wash Buffer (+EDTA) per 1 million cells and were aliquoted into 0.5 ml protein low binding tubes containing the primary histone mark antibody (1:200 dilution for H3K4me1 and H3K4me3 and 1:100 for H3K9me3 and H3K27me3) diluted in 200 μl Wash Buffer (+EDTA). Cells were incubated overnight at 4 °C on a roller, before they were washed once with 500 μl Wash Buffer. In the case of double labeling experiments, cells were incubated with antibodies against H3K4me1 and H3K9me3 together at the same concentrations as for the single mark experiments. We sorted four plates incubated simultaneously with H3K4me1 and H3K9me3.

Afterwards cells were resuspended in 500 μl Wash Buffer containing PaMN (3 ng/ml) and Hoechst 34580 (5 μg/ml) and incubated for 1h at 4 °C on a roller.

Finally, cells were washed an additional 2 times with 500 μl Wash Buffer before passing it through a 70 μm cell strainer (Corning, 431751) and sorting G1 cells based on Hoechst staining on an Influx FACS machine into 384 well plates containing 5 μl sterile filtered mineral oil (Sigma Aldrich) per well, always leaving eight wells empty of cells as a negative control. For bone marrow we sorted with the help of the indicated surface marker stainings unenriched (only using G1 gate), lineage negative, and LSK cells into separate parts of 384 well plates. We sorted 28, 26, 18, and 26 plates for H3K4me1, H3K4me3, H3K27me3, and H3K9me3, respectively. Of these, three plates were sorted with nuclei for unenriched cells only, three for lineage negative only, and three for LSK cells only for H3K4me1, H3K4me3, and H3K9me3. For lineage negative only and LSK cells only, cell populations were enriched by FACS before nuclei isolation. Five plates were sorted for both unenriched and lineage negative together. All remaining plates included a mixture of unenriched, lineage negative, and LSK sorted cells. The following small volumes were distributed using a Nanodrop II system (Innovadyme) and plates were spun for 2 min at 4 °C and 2000g after each reagent addition.

#### Protein A-MN activation

100 nl of Wash Buffer (-protease inhibitor), containing 2 mM CaCl_2_, were added per well to induce Protein A-MN mediated chromatin digestion that was performed for 30 min in a PCR machine set at 4 °C. Afterwards the reaction was stopped by adding 100 nl of a stop solution containing 40 mM EGTA (chelates Ca^2+^ and stops MN, Thermo, 15425795), 1.5% NP40 and 10 nl 2 mg/ml proteinase K (Invitrogen, AM2548) and incubated for further 20 min at 4 °C. Chromatin is subsequently released and PaMN permanently destroyed by proteinase K digestion at 65 °C for 6h followed by 80 °C for 20 min to heat inactivate proteinase K. Afterwards plates can be stored at −80 °C until further processing.

#### Library preparation

DNA fragments are blunt ended by adding 150 nl of the following mix per well and incubating for 30 min at 37 °C followed by 20 min at 75 °C for enzyme inactivation:

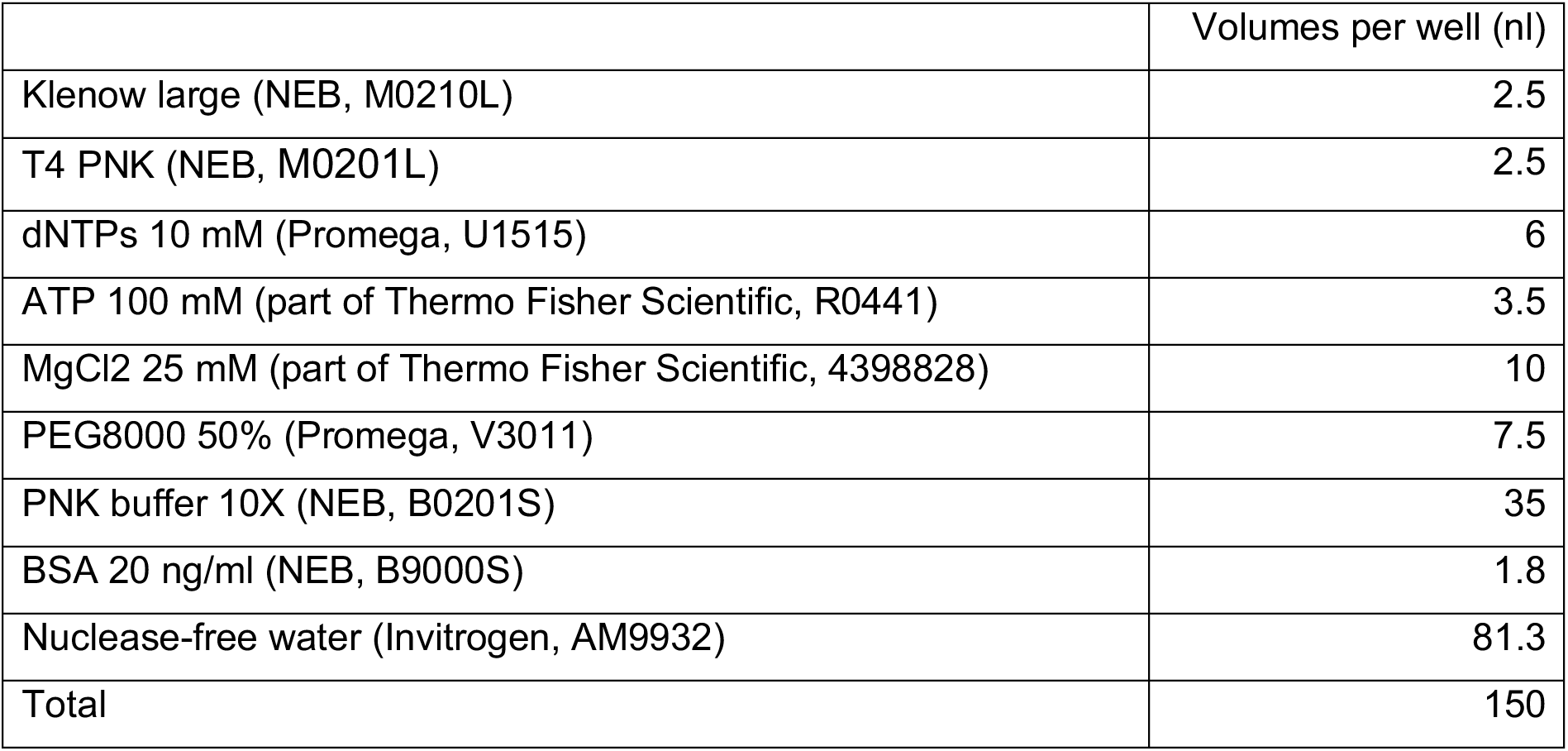

Blunt fragments are subsequently A-Tailed by adding 150 nl per well of the following mix and incubate for 15 min at 72 °C. Through AmpliTaq 360’s strong preference to incorporate dATP as a single base overhang even in the presence of other nucleotides, a general dNTP removal is not necessary:

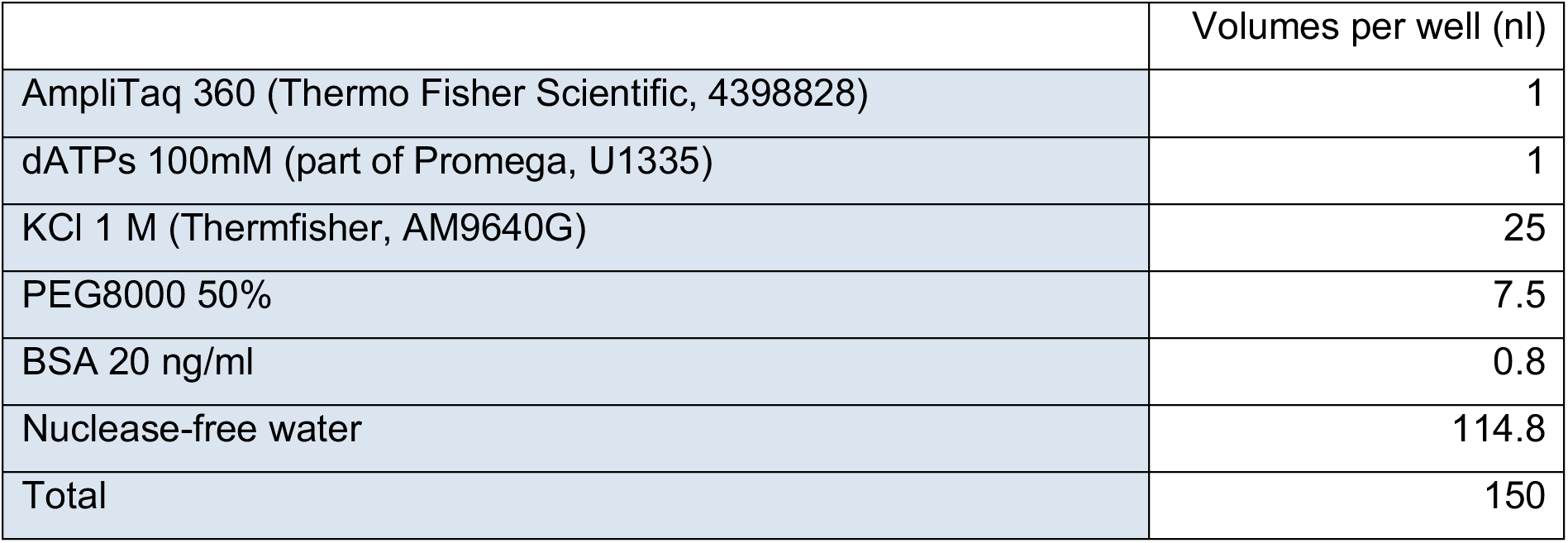

Next fragments are ligated to T-tail containing forked adaptors with following sequence: Top strand

**GGTGAT**GCCGGTAATACGACTCACTATAGGGAGTTCTACAGTCCGACGATC<u>NNN</u>*ACAC ACTA*T

Bottom strand

/5Phos/*TAGTGTGT*<u>NNN</u>GATCGTCGGACTGTAGAACTCCCTATAGTGAGTCGTATTACCG GC**GAGCTT**

Sequence features from left to right on the top strand:

Bases written in bold form a fork to prevent adaptor dimer- or multimerization. Bases in green represent T7 polymerase binding site for IVT based amplification. Bases in blue are the binding site (RA5) for the TruSeq Small RNA indexing primers (RPIx). The 3 random nucleotides underlined are the unique molecular identifier used for read deduplication and the 8 bases afterwards in italics represent the cell barcode which is different each of the 384 wells. For a full list of adaptors see Table S3.

For ligation 50 nl of 5 μM adaptor in 50 mM Tris pH7 is added to each well with a Mosquito HTS (ttp labtech). After centrifugation 150 nl of the following mix are added before plates are incubated for 20 min at 4 °C, followed by 16 h at 16 °C for ligation and 10 min at 65 °C to inactivate ligase:

|  | Volumes per well (nl) |
| --- | --- |
| T4 ligase (400K Units/ml, NEB, M0202L) | 25 |
| MgCl<sub>2</sub> 1 M (ThermoFisher, AM9530G) | 3.5 |
| Tris 1 M pH 7.5 (ThermoFisher, 15567027) | 10.5 |
| DTT 0.1M (Invitrogen, 15846582) | 52.5 |
| ATP 100 mM | 3.5 |
| PEG8000 50% | 10 |
| BSA 20 ng/ml | 1 |
| Nuclease-free water | 44 |
| Total | 150 |

Ligation products were pooled by centrifugation into oil coated lids of pipettip boxes at 200g for 2 min and the liquid face was transferred into 1.5 ml eependorf tubes and was purified by centrifugation at 13000g for 1 min and transfer into a fresh tube twice. DNA fragments were purified using Ampure XP beads (Beckman Coulter - prediluted 1 in 8 in bead binding buffer – 1 M NaCl, 20% PEG8000, 20 mM TRIS pH=8, 1 mM EDTA) at a bead to sample ratio of 0.8. Beads were washed twice with 1 ml 80% ethanol resuspending the beads during the first wash and resuspended in 8 μl Nuclease-free water and transferred into a fresh 0.5 ml tube. The cleaned DNA is then linear amplified by invitro transcription adding 12 μl of MEGAscript™ T7 Transcription Kit (Fishert Sc, AMB13345) for 12 h at 37 °C. Template DNA is removed by addition of 2 μl TurboDNAse (IVT kit) and incubation for 15 min at 37 °C. The produced RNA is further purified using RNA Clean XP beads (Beckman Coulter) at 0.8 beads to sample ratio, followed by RNA fragmentation for 2 min at 94 °C. After another bead cleanup, 40% (5 μl) of the RNA is primed for reverse transcription by adding 0.5 μl dNTPs (10 mM) and 1 μl randomhexamerRT primer 20 μM (GCCTTGGCACCCGAGAATTCCANNNNNN) and hybridizing it by incubation at 65 °C for 5 min followed by direct cool down on ice. Reverse transcription is performed by further addition of 2 μl first strand buffer (part of Invitrogen, 18064014), 1 μl DTT 0.1M (Invitrogen, 15846582), 0.5 μl RNAseOUT (Invitrogen, LS10777019) and 0.5 μl SuperscriptII (Invitrogen, 18064014) and incubating the mixture at 25 °C for 10 min followed by 1 h at 42 °C. Single stranded DNA is purified through incubation with μl RNAseA (Thermo Fisher, EN0531) for 30 min at 37 °C and PCR amplification to add the Illumina smallRNA barcodes and handles by adding 25 μl of NEBNext Ultra II Q5 Master Mix (NEB, M0492L), 11 μl Nuclease free water and 2 μl of RP1 and RPIx primers (10 μM). PCR cycles depended on the abundance of the histone modification assayed (8-10 for H3K9me3 and H3K27me3 10-12 for H3K4me1 and H3K4me3). Abundance and quality of the final library are assessed by QUBIT and bioanalyzer.

### Data preprocessing

Fastq files were demultiplexed by matching to an 8 nt cell barcode found in read 1 (R1). The 3 nt UMI was placed into the fastq header. To every read pair a MNase cut site is assigned to a genomic location. The cut site is defined as the genomic mapping location of the second base in R1. The ligation motif is defined as the two bases flanking the MNase cut site.

Assignment of read pairs to molecules is performed by pooling all read pairs that share the same UMI, cell barcode, and MNase cut site in a window of 1 kb.

We discarded read pairs if reads have:

- mapping quality scores (MAPQ) below 40,
- alternative hits at a non-alternative locus,
- mapped to separate locations beyond the expected insert size range,
- soft clips,
- more than 2 bases that differed from the reference,
- indels,
- mapping to a blacklist region (http://mitra.stanford.edu/kundaje/akundaje/release/blacklists/).

We selected cells with more than 500 total unique cuts for H3K4me1 and H3K4me3, and more than 1000 total unique cuts for H3K27me and H3K9me3. Cells also needed to have more than 50% of their cuts occur in an “AT” context. We also counted cut fragments that map in 50 kb nonoverlapping bins genome-wide, and calculated the fraction of bins that contains exactly zero cuts. Cells with a small fraction of zero cuts relative to other cells are more likely to have unspecific cuts. For each mark, we removed cells with a fraction of zero cuts that was below 2 standard deviations from the mean across all cells.

More details on the preprocessing pipeline can be found in the wiki page pipeline: https://github.com/BuysDB/SingleCellMultiOmics/wiki.

### Calculating reads falling in peaks in sortChIC for K562 cells

For each histone modification, we merged K562 single-cell sortChIC data, and used the resulting pseudobulk as input for *hiddenDomains* (Starmer and Magnuson, 2016), with minimum peak length of 1000 bp. We estimated 40574, 58257, 28499, and 28380 peaks for H3K4me1, H3K4me3, H3K27me3, and H3K9me3, respectively. For each histone modification, we counted the fraction of total reads that fall within each set of peaks.

### Dimensionality reduction based on multinomial models

We counted the number of cuts mapped to peaks across cells and applied the Latent Dirichlet allocation (LDA) model (Blei et al., 2003), which is a matrix factorization method that models discrete counts across predefined regions as a multinomial mixture model. LDA can be thought of as a discrete version of principal component analysis (PCA), replacing the normal likelihood with a multinomial one (Buntine and Jakulin, 2012). LDA models the genomic distribution of cuts from a single cell using a hierarchy of multinomials:

1. 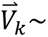 Dirichlet(*δ*) to specify the distribution over genomic regions for each topic *k* (length G genomic regions).
2. 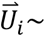 Dirichlet(*α*) to specify the distribution over topics for a cell *i* (length K topics). To generate the genomic location of the *j*th read in cell *i*:
3. Choose a topic 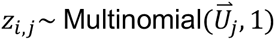
4. Choose a genomic region 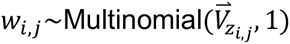

We used the LDA model implemented by the *topicmodels* R package (Grün and Hornik, 2011), to infer the cell-to-topic matrix (analogous to the scores matrix in PCA) and topic-to-region matrix (analogous to the loadings matrix in PCA) using Gibbs sampling with hyperparameters *α* =50/K, *δ* =0.1, where *K* is the number of topics. We used K=30 topics for all of our analyses.

### Defining eight sets of blood cell type-specific genes for cell typing

We defined cell type-specific genes for cell type calling by counting reads at +/− 5 kb centered at annotated transcription start sites (TSS). We applied LDA to the resulting count matrix for H3K4me1, H3K4me3, and H3K27me3 (we ignored H3K9me3 here because H3K9me3 marks mostly AT-rich, gene-poor regions). We found eight topics that defined the eight cell types in the data. For each topic, we took the top 150 TSS loadings to make eight sets of genes defining the cell types in the data. To compare this set of TSSs to publicly available scRNA-seq data (Giladi et al., 2018), we took each TSS and assigned it to the corresponding gene.

### Defining genomic regions for dimensionality reduction

We initially defined regions based on 50 kb windows genome wide, applying LDA, and using the Louvain method to define clusters to merge single-cell bam files. These merged bam files were then used to call significantly marked regions using *hiddenDomains* (Starmer and Magnuson, 2016) with minimum bin size of 1 kb. We merged the regions across clusters and generated a new count matrix using the *hiddenDomains* peaks as features. This new count matrix was used as input for dimensionality reduction.

### Batch correction in dimensionality reduction

Initial LDA of the count matrix revealed batch effects in H3K4me1 and H3K9me3 between cell types of plates that contained only one sorted type (i.e., entire plate was either unenriched, lineage-negative, or LSK cells, referred to as “single-type”) and cell types from plates that contained a mixture of unenriched and non-mature cells (referred to as “balanced”). We corrected batch effects in H3K4me1, H3K4me3, and H3K9me3. Since H3K27me3 did not have single-type plates, we did not correct batch effects in H3K27me3. We considered balanced plates as the reference for differences between cell types, and corrected deviations in single-type plates to match the balanced plates. We used the imputed sortChIC-seq signal inferred from LDA as a denoised signal *Y* for each genomic region *g* for every cell *c*:

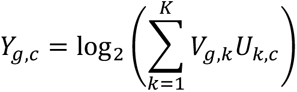

We modeled the cell type-specific batch effect using a linear model for each genomic region. The model infers the effect of a cell *c* belonging to batch *b* and cell type *d*:

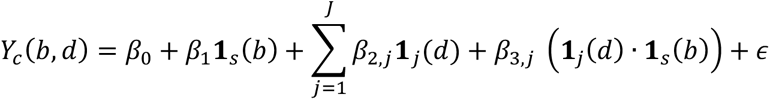

Where:

**1***_s_*(*b*) is an indicator variable equal to 1 if the cell is from a single-type plate (batch *s*), otherwise 0.

**1***_j_*(*d*) is equal to 1 if cell belongs to cell type *j*, otherwise 0.

*β*_0_ is the intercept of the model.

*β*_1_ is the global effect (i.e., independent of cell type) from a cell being from batch *s* (single-type plate).

*β*_2,*j*_ is the effect from a cell belonging to cell type *j*.

*β*_3,*j*_ is the interaction effect from a cell belonging to cell type *j* and being from batch *s* (single-type plate).

*∊* is Gaussian noise.

We inferred the effects for each genomic region using lm() in R with the formula syntax:

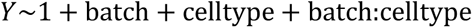

and estimated the batch-corrected signal:

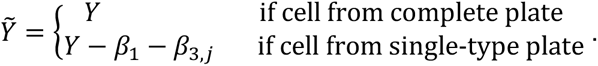

For cells that belong to complete plates, **1***_s_*(*b*) = 0. Therefore, this batch-correction only corrects signal from cells belonging to single-type plates. In the bone marrow analysis this corresponds to nine plates for H3K4me1, H3K4me3, and H3K9me3.

The corrected signal is used to refine the cell-to-topic and topic-to-region matrix by GLM-PCA (Townes et al., 2019). We applied SVD to the batch-corrected matrix to use as initializations for U and V, and included batch ID as cell-specific covariates (in glmpca R package: glmpca(), with fam=”poi”, minibatch=”stochastic”, optimizer=”avagrad”, niterations = 500). The batch-corrected U matrix is then visualized with uniform manifold approximation and projection (UMAP).

### Differential histone mark levels analysis

To calculate the fold change in histone mark levels at a genomic region between a cell type versus HSPCs, we modeled the discrete counts *Y* across cells as a Poisson regression. We fitted a null model, which is independent of cell type, and a full model, which depends on the cell type and compared their deviances to predict whether a region was “changing” or “dynamic” across cell types. We used the glm() implementation in R with the formula syntax for the full and null model:

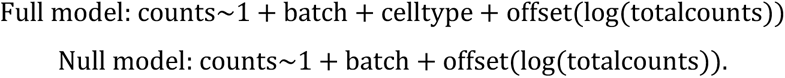

We used *G* as a deviance test statistic:

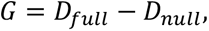

where the deviance is two times the log-likelihood, which for Poisson is:

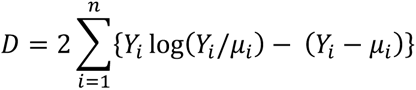

For the full model, the logarithm of the expected value *µ* is:

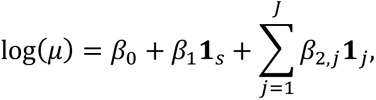

While for the null model, it is:

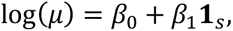

We fitted the model such that the estimated log_2_ relative to HSPCs. fold change of a cell type *j*, 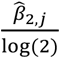, is always relative to HSPCs.

Under the null hypothesis, *G* is chi-squared distributed with degrees of freedom equal to the difference in the number of parameters in the two models. We use this test statistic to estimate a p-value and infer whether a 50kb bin is “changing” or “dynamic” across cell types. For H3K4me1, H3K4me3, and H3K27me3, we used a Benjamini-Hochberg adjusted p-value of q<10^−50^. For H3K9me3, where fold changes were generally smaller, we used q<10^−9^. This separate cutoff for H3K9me3 allowed comparable number of differential bins for downstream analysis.

### Defining bins above background levels for each mark

For each mark, we counted fragments falling in 50 kb bins summed across all cells. We then plotted this vector of summed counts as a histogram in log scale, which shows a bimodal distribution. We manually defined a cutoff for each mark as a background level, and took bins that were above this cutoff. This cutoff resulted in 22067, 12661, 18512, and 19881 bins for H3K4me1, H3K4me3, H3K27me3, and H3K9me3 respectively.

### Calculating bins that change independent of cell type

We defined “changing bins” or “dynamic bins” using the deviance test statistic, detailed above in “Differential histone mark analysis”, using a q-value<10^−50^ for H3K4me1, H3K4me3, and H3K27me3, and q<10^−9^ for H3K9me3. We defined these bins to be changing in a cell fate-independent manner if the estimated cell type effect, 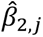, was either greater than 0 for all cell types (gained relative to HSPCs) or less than 0 for all cell types (lost relative to HSPCs).

### Predicting Activities of Transcription Factors in Single Cells

We adapted MARA (Motif Activity Response Analysis) described in (Arnold et al., 2013) to accommodate the sortChIC data. Briefly, we model the log imputed sortChIC-seq signal as a linear combination of TF binding sites and activities of TF motifs:

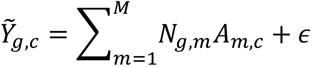

Where 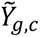 is the batch-corrected sortChIC-seq signal in genomic region *g* in cell *c*; *N_g,m_* is the number of TF binding sites in region *g* for TF motif *m*; *A_m,c_* is the activity of TF motif *m* in cell *c*; *∊* is Gaussian noise.

The single-cell motif activity, *A_m,c_*, is then overlaid onto the UMAP to show cell type-specific activities.

For H3K4me1, we defined genomic regions based on peak calling from *hiddenDomains*. For repressive marks, where domains can be larger, we used 50 kb bins that were significantly changing across cell types as genomic regions.

#### Creating the TF binding site matrix

We predicted the TF binding site count occurrence under each peak using the mm10 Swiss Regulon database of 680 motifs (http://swissregulon.unibas.ch/sr/downloads). We used the Motevo method to predict transcription factor binding sites. Posterior probabilities < 0.1 are rounded down to zero.

### Joint H3K4me1 and H3K9me3 analysis by double incubation

To simultaneously infer the H3K4me1 and H3K9me3 cluster from single-cell double-incubated cuts, we focused on regions that were most informative to distinguish between clusters in H3K4me1 and in H3K9me3. For H3K9me3, we used 6085 statistically significant changing bins (q<10^−9^, Poisson regression). For H3K4me1, we used regions near cell type-specific genes that were used to determine cell types from the data (811 regions). Since H3K4me1 had strong signal at both the TSS and gene bodies, we defined regions for each gene from transcription start site (TSS) to either its end site or 50 kb downstream of the TSS, whichever is smaller. We counted cuts mapped to these 6896 regions for H3K4me1, H3K9me3, as well as H3K4me1+H3K9me3 cells.

For a single cell, we assumed that the vector of H3K4me1+H3K9me3 counts 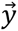 was generated by drawing *N* reads from a mixture of two multinomials, one from a cell type *c* from H3K4me1 (parametrized by relative frequencies 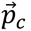 and one from a lineage *l* from H3K9me3 (parametrized by relative frequencies 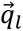):

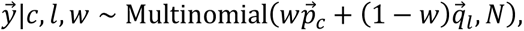

where *w* is the fraction of H3K4me1 that was mixed with H3K9me3.

Genomic region probabilities 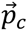 and 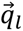 were inferred by the single-incubated data by averaging the imputed signal across cell types:

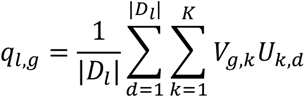

where *D_l_* is the set of cells that belong to lineage *l*. *V* and *U* are estimated from LDA.

The log-likelihood for the H3K4me1+H3K9me3 counts coming from cluster pair (*c, l*), can be defined as:

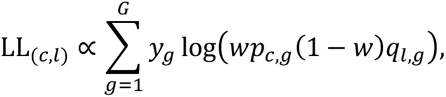

where g is a genomic region.

To assign a cluster pair to a double-incubated single cell, we calculated the log-likelihood for each possible pair (we had four lineages from H3K9me3 and eight clusters from H3K4me1, creating a 32 possible pairs) and selected the pair with the highest log-likelihood. We used the Brent method implemented in R (*optim*) to infer *w* that maximizes the log-likelihood for each pair.

### Materials

#### Antibodies

H3K4me1, ab8895 (Abcam), Lot: GR3206285-1

H3K4me3, 07-473 (Merck), Lot: 3093304

H3K9me3, ab8898 (Abcam), Lot: GR3217826-1

H3K27me3, 9733S (NEB), monoclonal

Public K562 data

H3K4me1, Peggy Farnham, ENCSR000EWC, pAb-037-050 (Diagenode)

H3K4me3, Peggy Farnheim, ENCSR000EWA, 9751S (Cell Signaling)

H3K9me3, Bradley Bernstein, ENCSR000APE, ab8898 (Abcam)

H3K27me3, Peggy Farnheim, ENCSR000EWB, 9733S (Cell Signaling)

Public bone marrow scRNA-seq:

Pseudobulk estimates merged from scRNA-seq data from (Giladi et al., 2018)

