## Supplemental Information for "Hierarchical chromatin regulation during blood formation uncovered by single-cell sortChIC"

**This PDF file includes:**

Figs. S1 to S6
Tables S2

**Other Supplementary Materials for this manuscript include the following:**

Tables S1 and S3

**Figure S1**

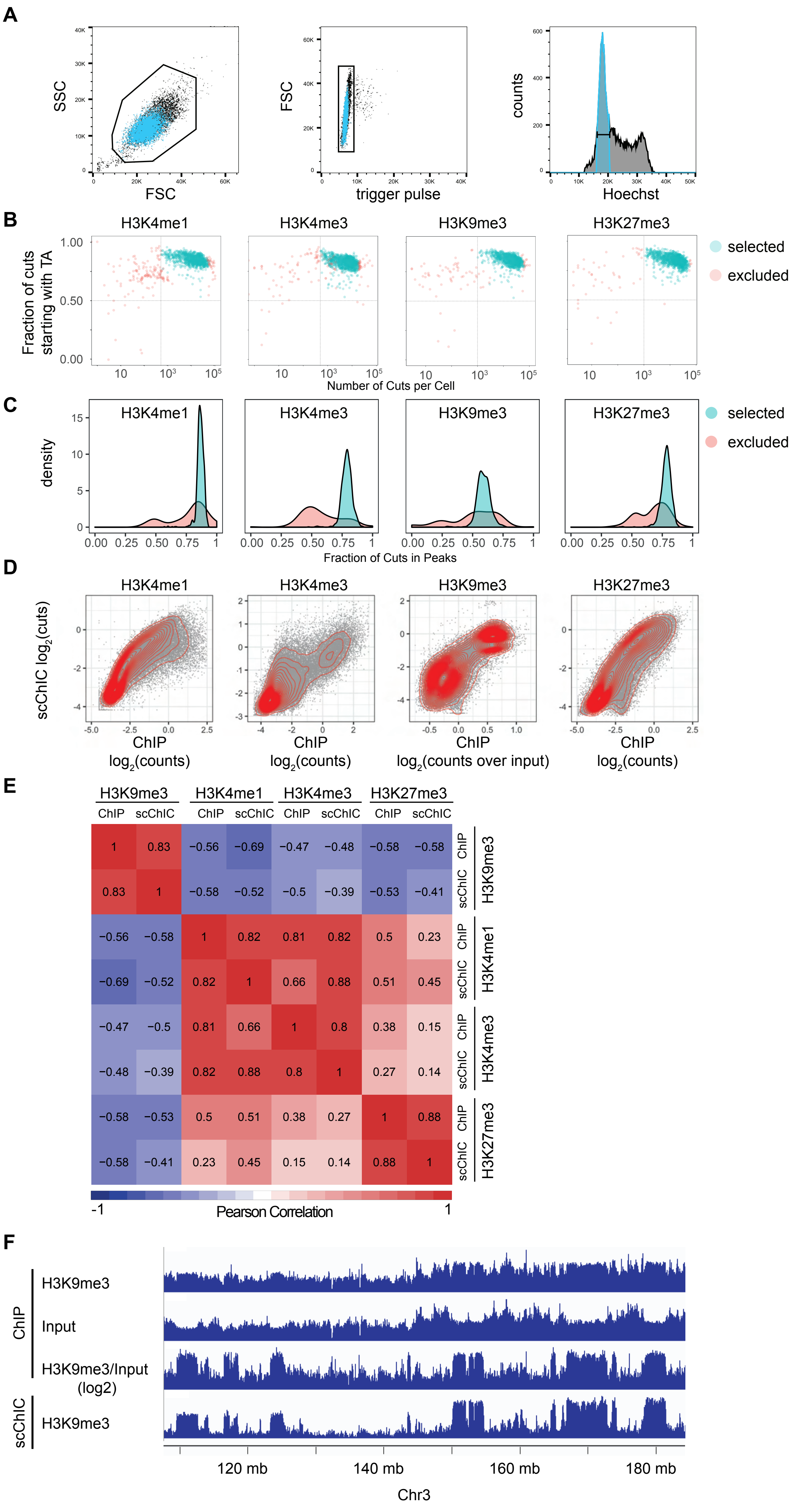
A
SSC
FSC
trigger pulse
counts
Hoechst
B
H3K4me1
H3K4me3
H3K9me3
H3K27me3
Fraction of cuts starting with TA
Number of Cuts per Cell
selected
excluded
C
density
Fraction of Cuts in Peaks
D
scChIC log2(cuts)
ChIP log2(counts)
ChIP log2(counts over input)
E
ChIP
scChIC
Pearson Correlation
F
ChIP
H3K9me3
Input
H3K9me3/Input (log2)
scChIC
120 mb
140 mb
160 mb
180 mb
Chr3

**Figure S1. sortChIC generates high-resolution maps of histone modifications in single cells**. (A) FACS plots for sorting individual K562 cells in G1 phase. (B) Fraction of cuts starting with TA (reflecting the preference of MNase to cut in an AT context) versus number of cuts mapped to the K562 genome. Cells below horizontal dotted lines and left of vertical lines are excluded from the analysis. (C) Distribution of fraction of cuts mapped to locations within peaks across cells. (D) Correlation between pseudobulk sortChIC and bulk ChIP signal using 50 kilobase (kb) bins for H3K4me1, H3K4me3, H3K27me3, and H3K9me3. (E) Pearson correlation between pseudobulk sortChIC and bulk ChIP signal using 50 kb bins across the four histone marks. (F) Three tracks of H3K9me3 ChIP-seq bulk data, one for H3K9me3 without normalization (H3K9me3), one for the input (Input), and one where H3K9me3 is normalized to the input (H3K9me3/input). Fourth track is H3K9me3 sortChIC pseudobulk, showing that H3K9me3 ChIP-seq requires normalizing by input to resemble sortChIC.

### Figure S2

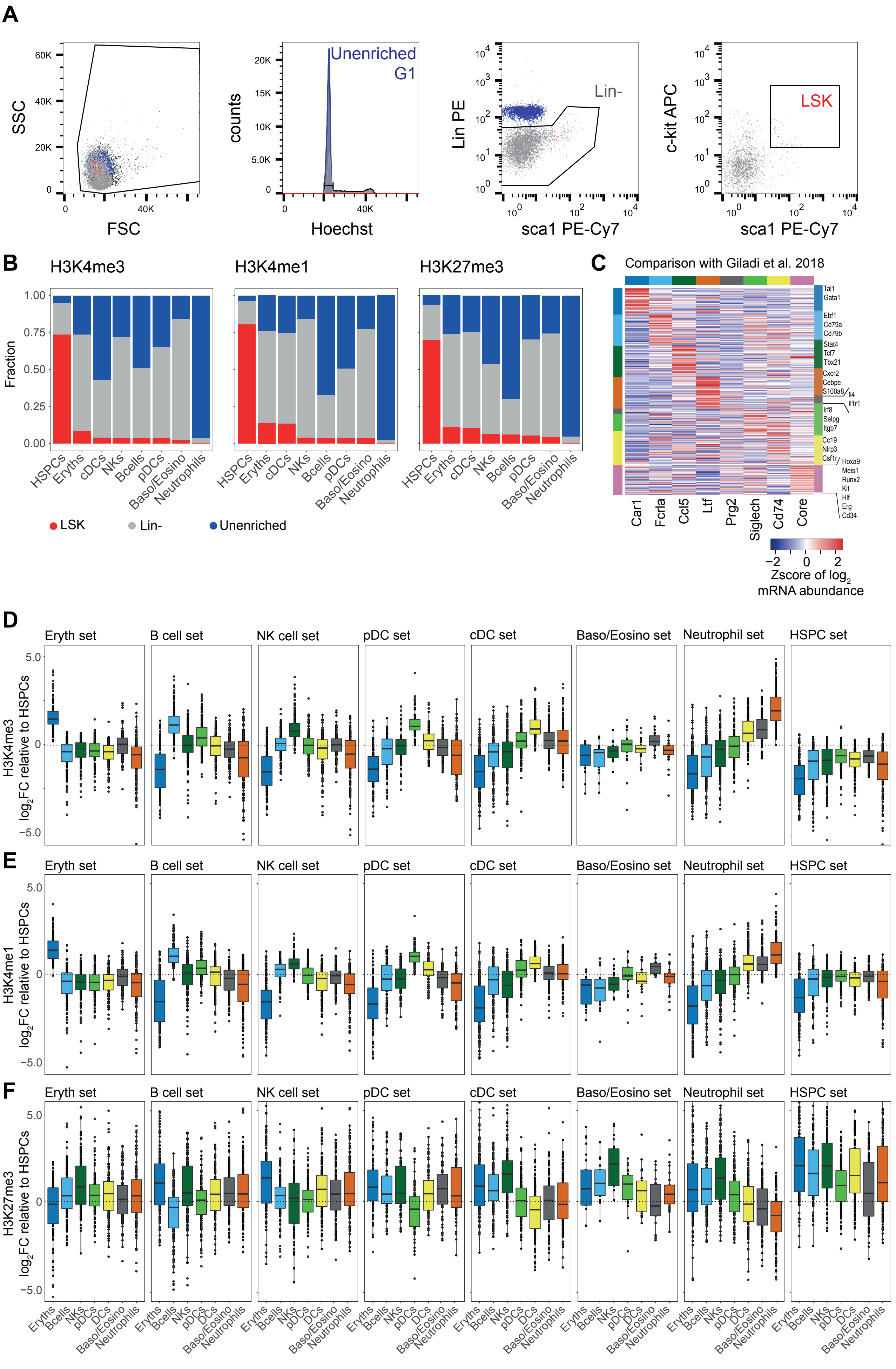
A
SSC
FSC
counts
Hoechst
Unenriched
G1
Lin PE
sca1 PE-Cy7
Lin-
c-kit APC
LSK
B
H3K4me3
H3K4me1
H3K27me3
Fraction
HSPCs
Eryths
cDCs
NKs
Bcells
pDCs
Baso/Eosino
Neutrophils
LSK
Lin-
Unenriched
C
Comparison with Giladi et al. 2018
Tal1
Gata1
Ebf1
Cd79a
Cd79b
Stat4
Tcf7
Tbx21
Cxcr2
Cebpe
S100a8
Il4
Il1r1
Irf8
Selpg
Itgb7
Cc19
Nlrp3
Csf1r
Hoxa9
Meis1
Runx2
Kit
Hlf
Erg
Cd34
Car1
Fcrla
Ccl5
Ltf
Prg2
Siglech
Cd74
Core
Zscore of log2 mRNA abundance
D
Eryth set
B cell set
NK cell set
pDC set
cDC set
Baso/Eosino set
Neutrophil set
HSPC set
H3K4me3
log2FC relative to HSPCs
E
H3K4me1
F
H3K27me3
Eryths
Bcells
NKs
pDCs
DCs
Baso/Eosino
Neutrophils

**Figure S2. H3K4me1 and H3K4me3 in HSPCs prime for different blood cell fates, while H3K27me3 in differentiated cell types silences genes of alternative cell fates**. (A) FACS plot for sorting G1 cells of whole bone marrow (unenriched), lineage negative ($Lin^-$), and $Lin^-$ ,$Sca1^+$, $cKit^+$ (LSK) populations. (B) Fraction of cells in each cell type labeled by the sorted population: whole bone marrow (unenriched), lineage negative ($Lin^-$), and $Lin^-Sca1^+cKit^+$ (LSK). (C) Cell type-specific mRNA abundances for genes associated with regions in Fig. 2E using pseudobulk analysis of the Giladi et al. 2018 dataset (Methods). (D) H3K4me3 fold changes of different cell types relative to HSPCs at cell type-specific regions. Each panel corresponds to a set of cell type-specific regions defined by the rows of one color in the heatmap of Fig. 2E. Regions are defined by +/- 5 kilobase windows centered at transcription start sites of cell type-specific genes. (E) Same as (D) but for H3K4me1. (F) Same as (D) but for H3K27me3.

#### Figure S3

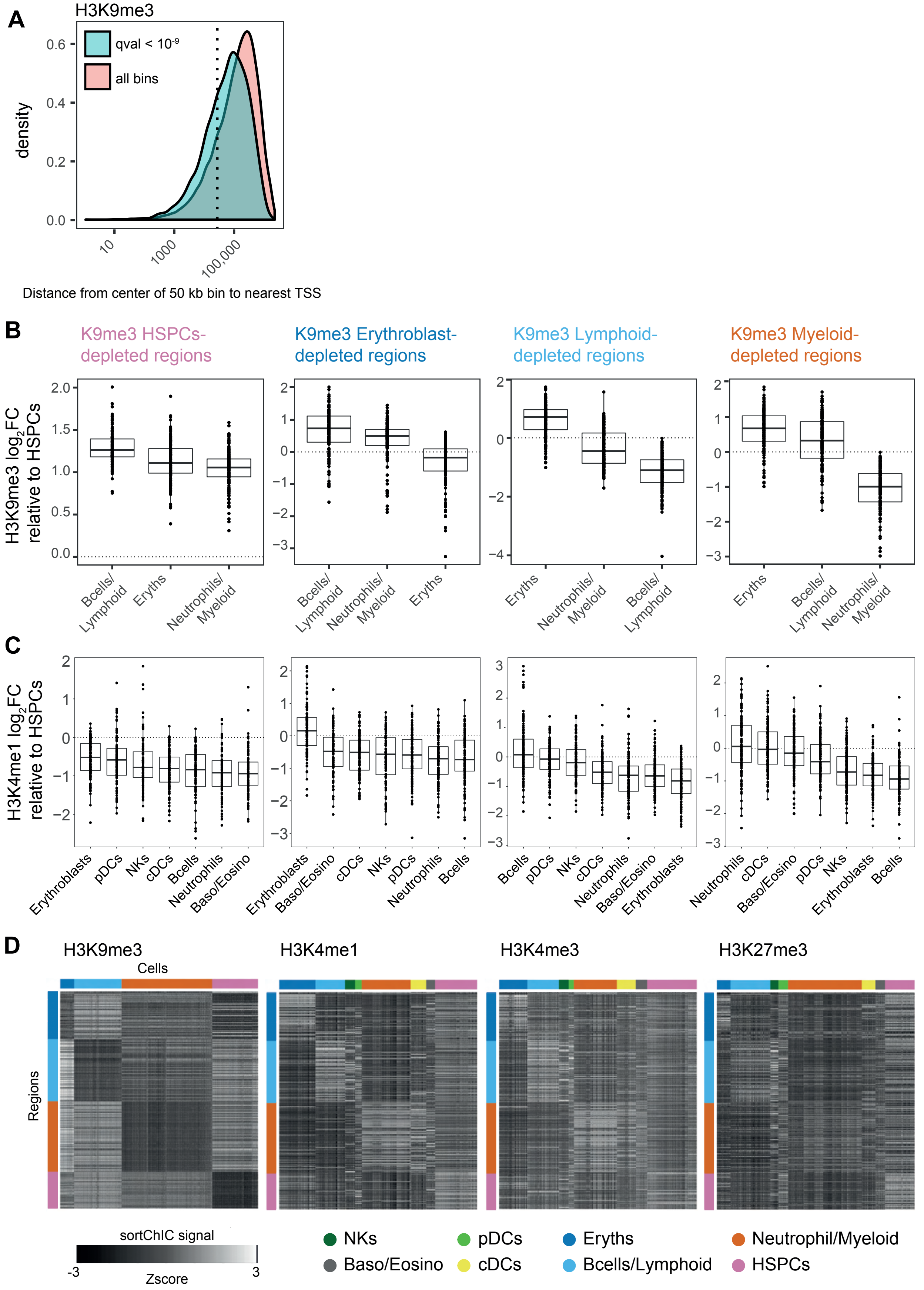
A
H3K9me3
qval < 10-9
all bins
density
0.6
0.4
0.2
0.0
10
1000
100,000
Distance from center of 50 kb bin to nearest TSS
B
K9me3 HSPCs-depleted regions
K9me3 Erythroblast-depleted regions
K9me3 Lymphoid-depleted regions
K9me3 Myeloid-depleted regions
H3K9me3 log2FC relative to HSPCs
Bcells/Lymphoid
Eryths
Neutrophils/Myeloid
C
H3K4me1 log2FC relative to HSPCs
Erythroblasts
pDCs
NKs
cDCs
Bcells
Neutrophils
Baso/Eosino
D
H3K9me3
H3K4me1
H3K4me3
H3K27me3
Cells
Regions
sortChIC signal
-3
Zscore
3
NKs
pDCs
Eryths
Neutrophil/Myeloid
Baso/Eosino
cDCs
Bcells/Lymphoid
HSPCs

**Figure S3. Lineage-specific loss of H3K9me3 correlates with cell type-specific increase in H3K4me1.** (A) Statistically significant 50 kb regions (adjusted p-value < $10^9$, deviance goodness-of-fit test) identified for H3K9me3, showing distribution of distances from center of 50 kb region to nearest gene. All bins are identified as 50 kb regions that have pseudobulk (counts summed across all cells) signal above background levels (Methods). Dotted line represents 25 kb, meaning the bin would overlap with a TSS. (B) Fold change in H3K9me3 relative to HSPCs for four sets of 150 regions: regions depleted in erythroblasts, lymphoid, myeloid, or HSPCs. Each region is 50 kb wide. (C) The same four sets of regions but showing fold change in H3K4me1, showing upregulation of H3K4me1 specifically in cell types that are depleted in H3K9me3. (D) Heatmap of the four regions in single cells across the four marks. Rows are regions, color coded as in top of (B). Columns are cells, color coded as in bottom.

### Figure S4

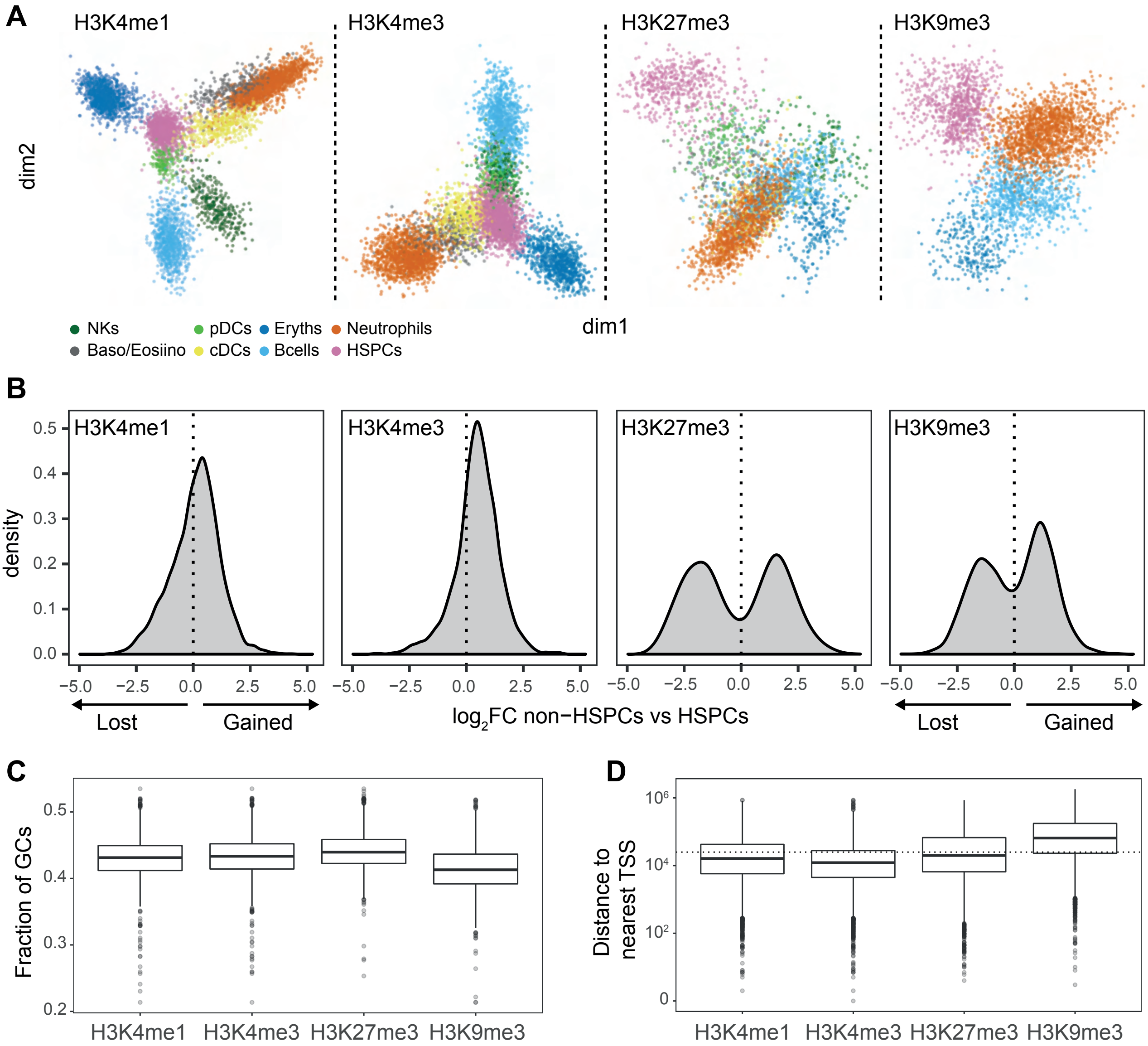


**E**

##### H3K9me3 regions unique to HSPCs

| GO biological process | GO number | P-value | Enrichment (log2) |
|---|---|---|---|
| complement activation, classical pathway | GO:0006958 | 5.68E-29 | 5.91 |
| phagocytosis, recognition | GO:0006910 | 1.86E-28 | 6.06 |
| B cell receptor signaling pathway | GO:0050853 | 3.58E-27 | 5.66 |
| phagocytosis, engulfment | GO:0006911 | 1.25E-26 | 5.46 |
| positive regulation of B cell activation | GO:0050871 | 4.69E-25 | 4.86 |
| defense response to bacterium | GO:0042742 | 2.33E-19 | 3.26 |
| innate immune response | GO:0045087 | 5.08E-11 | 2.24 |
| immunoglobulin production | GO:0002377 | 8.09E-08 | 3.13 |

##### H3K27me3 regions unique to HSPCs

| GO biological process | GO number | P-value | Enrichment (log2) |
|---|---|---|---|
| locomotory behavior | GO:0007626 | 9.29E-09 | 3.23 |
| central nervous system neuron differentiation | GO:0021953 | 5.14E-07 | 3.19 |
| regulation of nervous system process | GO:0031644 | 1.74E-06 | 3.31 |
| positive regulation of neuron differentiation | GO:0045666 | 2.10E-05 | 3.88 |
| developmental growth involved in morphogenesis | GO:0060560 | 4.64E-04 | 3.33 |
| neuromuscular process | GO:0050905 | 3.76E-04 | 3.28 |
| memory | GO:0007613 | 2.81E-04 | 3.19 |
| sensory perception of sound | GO:0007605 | 1.37E-04 | 3.16 |

**Figure S4. Features of active and repressive chromatin dynamics during hematopoiesis**. (A) Dimensionality reduction from GLMPCA (Methods) showing the two main latent factors explaining the sortChIC data for each mark. Dim 1 contains 8.2%, 8%, 7%, and 9.5% of the total L2 norm for H3K4me1, H3K4me3, H3K27me3, and H3K9me3, respectively. Dim 2 contains 7.9%, 6.8%, 6.8%, and 6.8% of the total L2 norm. (B) Distribution of $\log_2$ fold changes (FC) at statistically significant changing bins (null model: a bin has constant signal across all cell types, full model: a bin has signal that depends on cell type, deviance goodness-of-fit test) between pseudobulk of non-HSPCs versus HSPCs. Bimodal distribution highlights differences originate mainly between HSPCs and non-HSPCs. (C) GC content of dynamic 50 kb bins for the four histone marks. (D) Distance to nearest TSS measured from the center of each dynamic 50 kb bin. Dotted horizontal line represents 25 kb, meaning the bin would overlap with a TSS. (E) Gene ontology (GO) terms of HSPC-specific H3K9me3 (top) and H3K27me3 (bottom) regions. P-value and enrichment from Fisher's exact test.

**Figure S5**

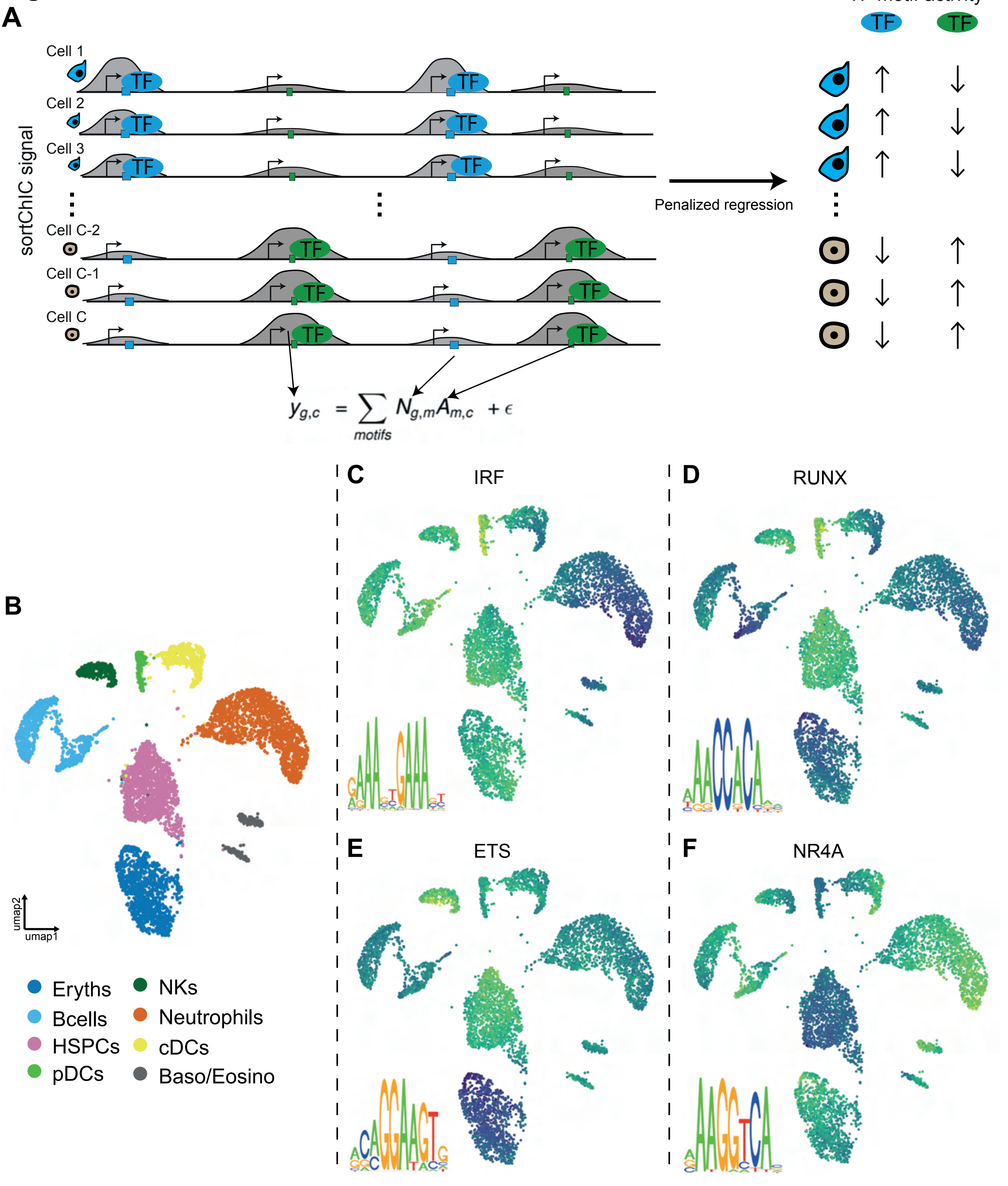
A
TF motif activity
TF
TF
Cell 1
Cell 2
Cell 3
Cell C-2
Cell C-1
Cell C
sortChIC signal
Penalized regression
$y_{g,c} = \sum_{motifs} N_{g,m} A_{m,c} + \epsilon$
B
C
IRF
D
RUNX
E
ETS
F
NR4A
umap1
umap2
Eryths
Bcells
HSPCs
pDCs
NKs
Neutrophils
cDCs
Baso/Eosino

**Figure S5. Penalized regression model reveals transcription factor motifs underlying cell type-specific chromatin dynamics.** (A) Schematic of the transcription factor (TF) activity model. The penalized regression model takes the imputed sortChIC signal in a peak as the response variable and the TFbinding motifs predicted under each peak as the explanatory variable (Method). The penalized multivariate regression infers the TF motif activity driving cell type-specific sortChIC signal. (B) UMAP of H3K4me1 chromatin states in single cells, colored by cell type. (C-F) UMAP where each cell is colored by the TF activity inferred from the model. Four cell type-specific TF motifs are shown.

### Figure S6

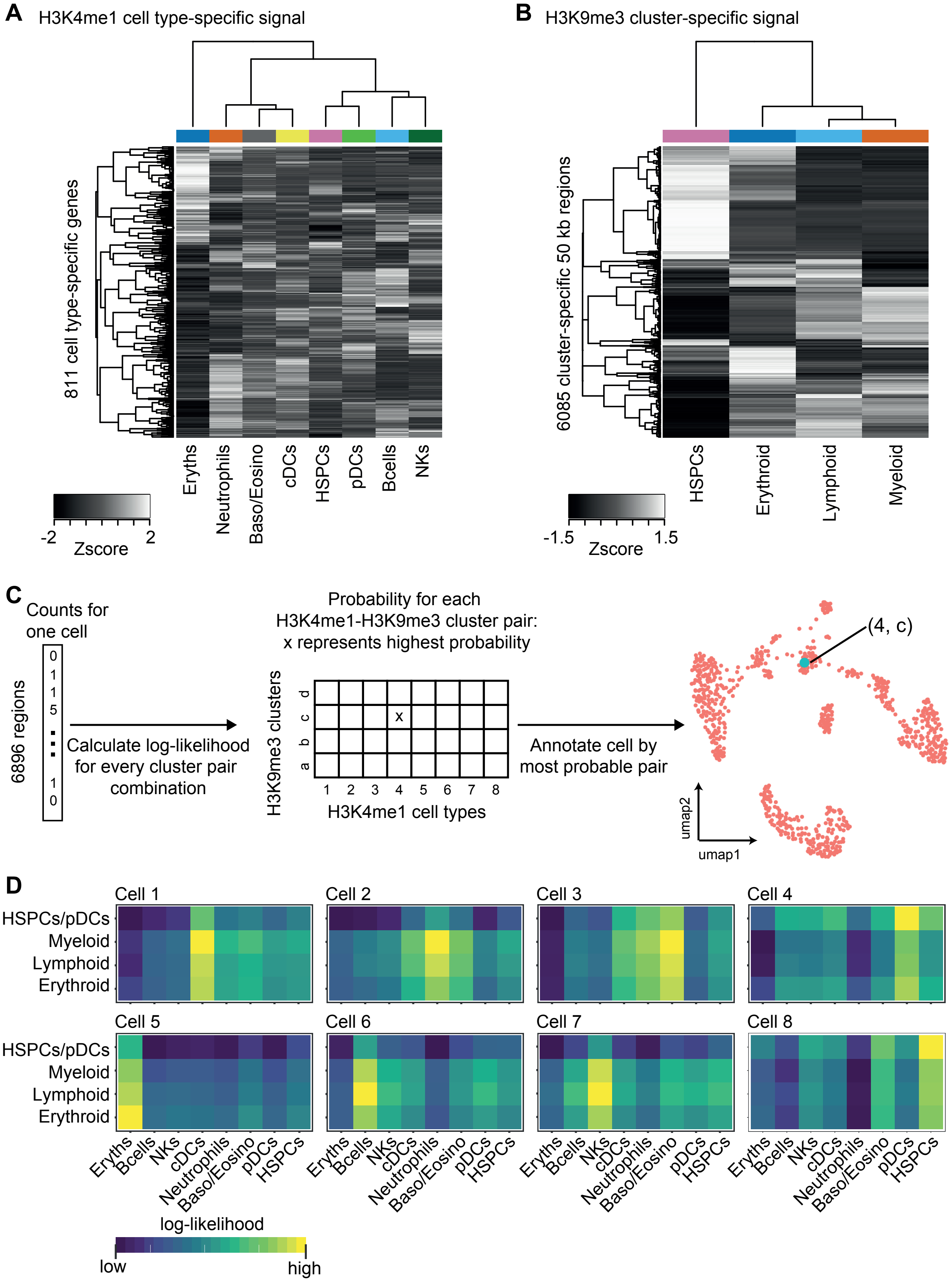

A
H3K4me1 cell type-specific signal
811 cell type-specific genes
Eryths
Neutrophils
Baso/Eosino
cDCs
HSPCs
pDCs
Bcells
NKs
-2
2
Zscore
B
H3K9me3 cluster-specific signal
6085 cluster-specific 50 kb regions
HSPCs
Erythroid
Lymphoid
Myeloid
-1.5
1.5
Zscore
C
Counts for one cell
6896 regions
0
1
1
5
1
0
Calculate log-likelihood for every cluster pair combination
Probability for each H3K4me1-H3K9me3 cluster pair: x represents highest probability
H3K9me3 clusters
d
c
b
a
x
1 2 3 4 5 6 7 8
H3K4me1 cell types
Annotate cell by most probable pair
(4, c)
umap2
umap1
D
Cell 1
Cell 2
Cell 3
Cell 4
Cell 5
Cell 6
Cell 7
Cell 8
HSPCs/pDCs
Myeloid
Lymphoid
Erythroid
Eryths
Bcells
NKs
cDCs
Neutrophils
Baso/Eosino
pDCs
HSPCs
log-likelihood
low
high

**Figure S6. Single-incubated data from H3K4me1 and H3K9me3 builds a model for inferring cluster-pairs in double-incubated data**. (A) Heatmap of H3K4me1 signal across clusters for 811 cell type-specific regions (Methods). These regions come from cell type-specific genes used in Figure 2E. (B) Heatmap of H3K9me3 signal across clusters for 6085 cluster-specific regions (50 kb genomic window). These regions come from the statistically significantly dynamic regions of H3K9me3 defined in Figure S3A. (C) Schematic of how a cluster-pair is inferred from each double-incubated cell. Each double-incubated cell has a vector of counts across 6896 regions (811 regions come from H3K4me1, while 6085 come from H3K9me3). We calculate the log-likelihood (Methods) of the observed double-incubated cell counts for each cluster-pair (32 cluster-pairs from 8 clusters in H3K4me1 and 4 clusters in H3K9me3). From the 32 log-likelihoods estimates, we assign the cell to the cluster-pair with the highest probability. (D) Examples of the 32 log-likelihood estimates from eight representative cells, shown as a 4-by-8 heatmap. Each of the four rows is a cluster from H3K9me3; each of the eight columns is a cluster from H3K4me1.

Table S2

| | ChIP based | | pA-MNase based | | | pA-Tn5 based | | | | | |
|---|---|---|---|---|---|---|---|---|---|---|---|
| Reference | Rotem et al. 2015 | Grosselin et al. 2019 | Hainer et al. 2019 | Ku et al. 2019 | This work | Harada et al. 2018 | Kaya-Okur et al. 2019 | Wang et al. 2019 | Bartosovik et al. 2021 | Janssens et al. 2020 | Wu et al. 2020 |
| Method | Sc-ChIP | High-throughput single-cell ChIP-seq | uliCUT&RUN | scChIC-seq | Sort-ChIC | ChIL-seq | CUT&Tag | CoBATCH | scCut&Tag | autoCUT&TAG | scCut&Tag |
| Integrated with FACS sorting | | | | | Hoechst, Lin, sca1,c-kit | | | | | | |
| Number of cells profiled (cell lines) | 7308 | 0 | 172 | 387 | 4136 | 15 | 1764 | 2161 | 8745 | NA | 2794 |
| Number of cells profiled (primary cells) | 0 | 7465 | 0 | 285 | 12208 | 0 | 0 | 3998 | 47340 | 6000 | 1311 |
| Approximate throughput (cells/run) | 100 | 1000 | low | low | 4500 (384 per plate) | low | 1000 | 2000 | 4000 | 2500 | 2794 |
| Approximate average number of reads/cell | 453-773 | 1630 | NA | 10000-15000 | 15000 | 15000-350000 | NA | 7525-12000 | 48-453 | 3900-13000 | 1729 |
| TFs and other non-histone proteins | | | CTCF, Nanog, Sox2 | | | | | Pol2 | Olig2, Rad21 | | |
| H3K4me1 (active) | X | | | | X | | | | | | |
| H3K4me2 (active) | X | X | | | | | X | | | | |
| H3K4me3 (active) | | | | X | X | X | | | X | X | |
| H3K36me3 (active) | | | | | | | | X | X | X | |
| H3K27ac (active) | | X | | | | X | | X | X | | |
| H3K27me3 (repressive) | | | | X | X | X | X | | X | X | X |
| H3K9me3 repressive) | | | | | X | | | | | | |

**Table S2. Comparison of studies on single-cell histone modification mapping.**
